# Neuroanatomical and Functional Consequences of Oxytocin Treatment at Birth

**DOI:** 10.1101/2022.05.21.492438

**Authors:** William M. Kenkel, Richard J. Ortiz, Jason R. Yee, Allison M. Perkeybile, Praveen Kulkarni, C. Sue Carter, Bruce S. Cushing, Craig F. Ferris

**Affiliations:** University of Delaware; Indiana University; University of Texas El Paso, New Mexico State University; University of Veterinary Medicine, Vienna; University of Virginia; Northeastern University; Kinsey Institute, Indiana University; University of Texas El Paso

## Abstract

Birth is a critical period for the developing brain, a time when surging hormone levels help prepare the fetal brain for the tremendous physiological changes it must accomplish upon entry into the ‘extrauterine world’. A number of obstetrical conditions warrant manipulations of these hormones at the time of birth, but we know little of their possible consequences on the developing brain. One of the most notable birth signaling hormones is oxytocin, which is administered to roughly 50% of laboring women in the United States prior to / during delivery. Previously, we found evidence for behavioral, epigenetic, and neuroendocrine consequences in adult prairie vole offspring following maternal oxytocin treatment immediately prior to birth. Here, we examined the neurodevelopmental consequences in adult prairie vole offspring following maternal oxytocin treatment immediately. Control prairie voles and those exposed to 0.25 mg/kg oxytocin were scanned as adults using anatomical and functional MRI, with neuroanatomy and brain function analyzed as voxel-based morphometry and resting state functional connectivity, respectively. Overall, anatomical differences brought on by oxytocin treatment, while widespread, were generally small, while differences in functional connectivity, particularly among oxytocin-exposed males, were larger. Analyses of functional connectivity based in graph theory revealed that oxytocin-exposed males in particular showed markedly increased connectivity throughout the brain and across several parameters, including closeness and degree. These results are interpreted in the context of the organizational effects of oxytocin exposure in early life and these findings add to a growing literature on how the perinatal brain is sensitive to hormonal manipulations at birth.

## BACKGROUND

Oxytocin (OXT) is a potent and pleiotropic hormone that surges at birth to help facilitate the tremendous changes that both mammalian mothers and their offspring must accomplish upon delivery (1–3). As of 2019, 29.4% of laboring women in the U.S. received OXT to induce labor (4), and according to available survey data, this figure raises to ∼50% of birthing women in America when considering OXT used to either induce and/or augment labor (5). This obstetric practice is of interest to neuroscience because there is evidence OXT can cross the placenta (6) and a growing literature suggests the neonatal brain is particularly sensitive to OXT around the time of birth, when OXT receptor (*Oxtr*) expression begins to accelerate (7) and OXT neurons in the brain undergo intense remodeling (8). Indeed, the long-term, developmental effects of OXT manipulations in early life are well-documented (9,10), which suggests the perinatal period may be a *sensitive period* with regard to the impact of OXT.

Some initial studies suggested higher rates of autism spectrum and attention deficit / hyperactivity disorder amongst children born to women whose labors were induced with OXT; however, meta-analysis of these findings suggest that any such conclusions remain premature (11). While there have been conflicting reports as to whether OXT administered to induce / augment labor is associated with increased rates of autism spectrum disorder or autistic-like behavior in offspring, considerations of dose add an important degree of nuance to this topic (12,13). Thus, regardless of whether the consequences of obstetrically administered OXT raise to the level of a neurodevelopmental disorder, the question of whether OXT affects offspring neurodevelopment is of great public health relevance given its widespread use.

Previously, we investigated the impact of maternally administered OXT on offspring neurodevelopment and behavior using the socially monogamous prairie vole (14). We found that fetal physiology was indeed sensitive to maternally administered OXT and that, in the fetal brain, such OXT dose-dependently increased methylation of the *Oxtr* promoter. In adulthood, OXT-exposed offspring of both sexes were found to demonstrate a broadly gregarious phenotype such that they exhibited more spontaneous alloparental care toward unrelated pups and spent more time in close social contact with opposite-sex adults. Male voles exposed to OXT also showed increased density of OXT receptor in the central amygdala, insular cortex, and parietal cortex, while showing decreased vasopressin receptor density in the ventral pallidum.

Because OXT is a pleiotropic hormone, we opted for a broad survey of the brain in the present study, carrying out whole-brain resting functional connectivity to further characterize the scope of the neurodevelopmental consequences of OXT exposure at birth. Here, we used magnetic resonance imaging (MRI) to scan the brains of adult male and female prairie vole offspring originally born to pregnant females treated with 0.25 mg/kg OXT on the expected day of delivery. We examined both anatomical measures (voxel based morphometry (VBM) and diffusion-weighted imaging (DWI)) as well as functional measures (resting state functional connectivity (rs-fMRI) and several graph theory measures detailed below).

## METHODS

### Subjects

Prairie vole offspring (*Microtus ochrogaster*) were generated as previously described (14). All procedures were conducted in accordance with the National Institutes of Health Guide for the Care and Use of Laboratory Animals and were approved by the Institutional Animal Care and Use Committee of Northeastern University. On the expected day of delivery, pregnant females were either injected intraperitoneally with OXT (0.25 mg/kg, ‘OXT’) or left undisturbed (‘Control’). Offspring were only included if they were delivered within 24 hours of OXT treatment. Offspring were raised by their birth parents as we previously observed no effect of maternal-OXT treatment on offspring outcomes (14). At 20 days of age, OXT and Control offspring were weaned into same-sex sibling pairs and were left to mature. Upon reaching adulthood (postnatal days 60-70), OXT and Control offspring underwent three neuroimaging scans: aT1-weighted anatomical scan for voxel based morphometry scan (VBM), an awake, resting state functional scan (rs-fMRI), and an anesthetized diffusion-weighted imaging scan (DWI), as detailed below. Subject offspring consisted of 17 Control females, 19 Control males, 17 OXT females, and 16 OXT males. From these, 7 Control females, 12 Control males, 13 OXT females and 13 OXT males were ultimately included in the rs-fMRI analyses after removing subjects due to motion artefact or technical difficulties.

### Neuroimaging

All neuroimaging measures were collected using a Bruker BioSpec 7.0T/20-cm Ultra Shield Refrigerated horizontal magnet (Bruker, Billerica, MA). A 20-G/cm magnetic field gradient insert (inner diameter 12 cm) was used to scan anesthetized subjects using a quadrature transmit/receive volume coil (inner diameter 38 mm). Imaging sessions began with an anatomical scan with the following parameters: 20 slices; slice thickness, 0.70 mm; field of view, 2.5 cm; data matrix, 256 × 3 × 256; repetition time, 2.5 seconds; echo time (TE), 12.0 ms; effective TE, 48 ms; number of excitations, 2; and total acquisition time, 80 seconds.

### Voxel Based Morphometry (VBM)

The following procedures were adapted for use in the vole from those described previously for rats (15). For each subject, the atlas (image size 256 × 256 × 63) (H × W × D) was warped from the standard space into the subject image space (image size 256 × 256 × 40) using the nearest-neighbor interpolation method. In the volumetric analysis, each brain region was therefore segmented, and the volume values were extracted for all 111 regions of interest (ROIs), calculated by multiplying unit volume of voxel (in mm^3^) by the number of voxels using an in-house MATLAB script. To account for different brain sizes, all ROI volumes were normalized by dividing each subject’s ROI volume by their total brain volume.

### Diffusion-weighted Imaging (DWI)

The following procedures were identical to those described previously (16,17). Diffusion-weighted imaging (DWI) was acquired with a spin-echo echo-planar imaging (EPI) pulse sequence with the following parameters: repetition time/TE, 500/20 ms; 8 EPI segments; and 10 noncollinear gradient directions with a single b-value shell at 1000 seconds/mm^2^ and 1 image with a b-value of 0 seconds/mm^2^ (referred to as b0). Geometrical parameters were as follows: 48 coronal slices, each 0.313 mm thick (brain volume) and with in-plane resolution of 0.313 × 3 × 0.313 mm^2^ (matrix size, 96 × 3 × 96; field of view, 30 mm^2^). The imaging protocol was repeated 2 times for signal averaging. DWI acquisition took 35 to 70 minutes. DWI included diffusion-weighted three-dimensional EPI image analysis producing fractional anisotropy (FA) maps and apparent diffusion coefficient. DWI analysis was implemented with MATLAB (version 2017b) (The MathWorks, Inc., Natick, MA) and MedINRIA version 1.9.0 (http://www-sop.inria.fr/asclepios/software/MedINRIA/index.php) software.

Each brain volume was registered with the three-dimensional MRI Vole Brain Atlas template (Ekam Solutions LLC, Boston, MA) allowing voxel- and region-based statistics (18). In-house MIVA software was used for image transformations and statistical analyses. For each vole, the b0 image was coregistered with the b0 template (using a 6-parameter rigid-body transformation). The coregistration parameters were then applied on the DWI indexed maps for each index of anisotropy. Normalization was performed on the maps providing the most detailed and accurate visualization of brain structures. Normalization parameters were then applied to all indexed maps and then smoothed with a 0.3-mm Gaussian kernel. To ensure that preprocessing did not significantly affect anisotropy values, the nearest neighbor option was used following registration and normalization.

### Resting State Functional MRI (rs-fMRI)

We used the same equipment and scanning protocols as in our recent work; for complete details see (18–20). Data were analyzed as 111 nodes corresponding to brain regions specified in a vole-specific atlas (18). Pearson’s correlation coefficients were computed per subject across all node pairs (6105), assessing temporal correlations between brain regions. Then, r-values’ (−1 to 1) normality were improved using Fisher’s Z-transform. For each group, 111×111 symmetric connectivity matrices were constructed, each entry representing the strength of edge. An |Z|=2.3 threshold was used to avoid spurious or weak node connections (21).

### Network Analyses

Graph theory network analysis was generated using Gephi, an open-source network and visualization software (22). For all groups, the absolute values of their respective symmetric connectivity matrices were imported as undirected networks and a threshold of |Z|=2.3 was applied to each node’s edges to avoid spurious or weak node connections (23).

### Betweenness Centrality

Betweenness centrality analyzes occurrences where a node lies in the path connecting other nodes (24). Let 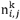 be the number of pathways from *i* to *j* going through *k*. Using these measures of connection, the betweeness of vertex *k* is:

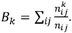

### Degree Centrality

Degree centrality indicates the number of associations of a specific node (25). Non-weighted, binary degree is defined as:

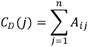

where *n* is the number of rows in the matrix in the adjacency matrix ***A*** and the elements of the matrix are given by *A*_*ij*_, the number of edges between nodes *i* and *j*.

### Closeness Centrality

Closeness centrality measures the average distance from a given starting node to all other nodes in the network (26). Closeness is defined as:

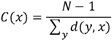

where *d(y,x*) is the distance between vertices *x* and *y* and *N* is the number of nodes in the graph.

### Statistics

Normality tests of control females, control males, OXT females and OXT males were performed to examine if parametric or non-parametric assumptions were required for future analysis. Shapiro-Wilk’s tests were performed to examine normality assumption for degree, closeness and betweeness centrality values. Regional p-values that were greater than 0.05 were assumed to be normal. A corresponding list of nodes that classified a region is detailed in table S1. After assumptions of normality were validated, one-way ANOVA tests were used to compare differences in degree, closeness and betweenness centralities between groups. When necessary, a nonparametric Kruskal-Wallis test was performed if there was evidence against normality assumption. Statistical differences between groups were determined using a Mann-Whitney U test (a = 5%). The following formula was used to account for false discovery from multiple comparisons:

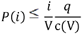

P(i) is the p value based on the t test analysis. Each of 111 regions of interest (ROIs) (i) within the brain containing V ROIs. For graph theory measures, statistical analyses were calculated using GraphPad Prism version 9.0.0 for MacOS (GraphPad Software, San Diego, California USA, www.graphpad.com). For post-hoc analyses of graph theory parameters, Holm-Šídák test (for parametric) and Dunn’s (for nonparametric) were used after correction for multiple comparisons.

## RESULTS

We observed a number of differences in VBM measures; however most were small to very small in effect size (Figure 1 and Tables 1-3). Within the Control group, there was only a single sex difference in regional volume; within the OXT group, however, OXT females had larger volumes in 8 of 17 cortical regions and smaller volumes in 11 brainstem / cerebellar regions compared to OXT males (Table 1). Similarly, Control males had larger volumes in 9 of 17 cortical regions and smaller volumes in 9 brainstem / cerebellar regions compared to OXT males (Table 2). Comparing within females, OXT treatment at birth resulted in smaller volumes in 4 of 17 cortical regions (Table 3). Control animals generally had larger amygdalar volumes than OXT animals (4 of 6 subregions in males; 2 of 6 in females). When morphometry data from all 111 brain regions were loaded into a PCA, the overall explanatory value of dimensions 1 and 2 was modest (29.4% and 15.9% respectively) and there were impacts of both sex and treatment on dimension 1, with male sex (F(1,69) = 4.98, p = 0.029) and OXT treatment (F(1,69) = 16.06, p < 0.001) leading to greater values (Figure 2).

**Table 1.** A) a list of the single brain area, reticular nucleus of the thalamus, that significantly differs (p=0.021, critical value p<0.05) in volume between adult female and male voles that were treated with saline vehicle within 24 hrs of parturition. Shown are the average and standard deviation in volume in mm3 and effects size (omega square ωSq). With a false discovery rate (FDR) p=0.0017 this singular finding can be dismissed concluding there is no sex difference in brain volumes between female and male voles resulting from this early manipulation. In contrast, Table 1B lists the brain areas that are significantly different in volume between adult female and male voles exposed to OXT during birth (FDR p=0.043). The brain areas are ranked in order of significance and are truncated from a larger list of 116 areas taken from the vole MRI atlas (see Supplementary Table S1).

| A | Female Vehicle vs Male Vehicle - Brain Volumes (mm3) |  |  |  |  |  |  |  |
| --- | --- | --- | --- | --- | --- | --- | --- | --- |
|  | Brain Area | Female |  |  | Male |  | P-val | ω Sq |
|  |  | Ave | SD |  | Ave | SD |  |  |
|  | reticular n. | 2.6 | 0.4 | > | 2.3 | 0.3 | 0.021 | 0.16733 |

| B | Female Oxytocin vs Male Oxytocin - Brain Volumes (mm3) |  |  |  |  |  |  |  |
| --- | --- | --- | --- | --- | --- | --- | --- | --- |
|  | Brain Area | Female |  |  | Male |  | P-val | ω Sq |
|  |  | Ave | SD |  | Ave | SD |  |  |
|  | frontal association ctx | 12.4 | 3.1 | > | 9.7 | 2.8 | 0.003 | 0.16823 |
|  | ventral tegmental area | 0.6 | 0.2 | < | 0.8 | 0.3 | 0.008 | 0.1341 |
|  | vestibular n. | 1.5 | 0.6 | < | 2.2 | 0.8 | 0.009 | 0.12879 |
|  | agranular insular ctx | 11.6 | 2.7 | > | 9.7 | 2.5 | 0.010 | 0.12744 |
|  | retrosplenial ctx | 7.5 | 0.8 | > | 6.7 | 1.0 | 0.014 | 0.11251 |
|  | gigantocellular reticular n. | 5.1 | 1.3 | < | 5.8 | 1.6 | 0.017 | 0.10415 |
|  | median raphe n. | 0.5 | 0.2 | < | 0.8 | 0.4 | 0.019 | 0.10093 |
|  | secondary motor ctx | 13.1 | 2.2 | > | 11.4 | 2.5 | 0.019 | 0.10065 |
|  | anterior hypothalamus | 3.6 | 0.9 | > | 3.1 | 1.0 | 0.019 | 0.09949 |
|  | parasubiculum | 1.4 | 0.4 | < | 1.8 | 0.5 | 0.021 | 0.09613 |
|  | paraflocculus cerebellum | 5.4 | 1.7 | < | 6.1 | 1.6 | 0.022 | 0.09494 |
|  | cuneate n. | 0.1 | 0.2 | < | 0.4 | 0.5 | 0.026 | 0.08752 |
|  | parietal ctx | 2.5 | 0.7 | > | 2.1 | 0.6 | 0.029 | 0.08393 |
|  | 7th cerebellar lobule | 0.6 | 1.4 | < | 2.3 | 2.8 | 0.031 | 0.08128 |
|  | claustrum | 1.9 | 0.5 | > | 1.5 | 0.6 | 0.031 | 0.08077 |
|  | medullary reticular n. | 0.9 | 2.2 | < | 2.7 | 3.4 | 0.036 | 0.07577 |
|  | olivary n. | 1.1 | 0.6 | < | 1.5 | 0.7 | 0.036 | 0.07557 |
|  | rostral piriform ctx | 11.9 | 1.7 | > | 10.5 | 2.3 | 0.039 | 0.07242 |
|  | reticulotegmental n. | 1.9 | 0.7 | < | 2.3 | 0.6 | 0.041 | 0.07042 |
|  | pontine reticular n. oral | 2.0 | 0.8 | < | 2.7 | 1.2 | 0.042 | 0.06945 |
|  | anterior olfactory n. | 14.1 | 2.1 | > | 12.2 | 2.8 | 0.045 | 0.06742 |
|  | anterior cingulate ctx | 4.1 | 1.1 | > | 3.5 | 0.8 | 0.045 | 0.06742 |
|  | glomerular layer olfactory bulb | 16.6 | 2.9 | > | 15.2 | 2.9 | 0.046 | 0.06642 |
|  | lateral preoptic area | 1.4 | 0.3 | > | 1.2 | 0.4 | 0.052 | 0.06168 |
|  | caudal piriform ctx | 9.0 | 1.4 | > | 8.3 | 1.2 | 0.052 | 0.0616 |

**Table 2.** The list of brain regions that were significantly different in volume between OXT females and Control females. OXT females showed smaller brain volumes in 15/21 of the affected regions (FDR p=0.036). The regions affected spread across the olfactory system (anterior olfactory nuc., piriform cortex), hypothalamus (paraventricular, anterior), amygdala (basal, extended), thalamus (anterior, paraventricular) and basal ganglia (nuc. accumbens, caudate putamen).

| Female Vehicle vs Female Oxytocin - Brain Volumes (mm3) |  |  |  |  |  |  |  |
| --- | --- | --- | --- | --- | --- | --- | --- |
| Brain Area | Female Veh |  |  | Female OT |  | P-val | ω Sq |
|  | Ave | SD |  | Ave | SD |  |  |
| tegmental n. | 1.3 | 0.3 | > | 0.9 | 0.3 | 0.000 | 0.34584 |
| paraventricular hypothalamus | 0.7 | 0.2 | > | 0.4 | 0.2 | 0.001 | 0.28447 |
| anterior thalamus | 2.2 | 0.7 | > | 1.5 | 0.5 | 0.005 | 0.20755 |
| anterior hypothalamus | 4.5 | 0.9 | > | 3.6 | 0.9 | 0.007 | 0.18794 |
| accumbens shell | 4.5 | 1.5 | > | 3.3 | 1.0 | 0.009 | 0.17454 |
| caudate putamen (striatum) | 25.7 | 4.1 | > | 22.2 | 3.4 | 0.010 | 0.16929 |
| solitary tract n. | 0.2 | 0.3 | < | 0.5 | 0.4 | 0.012 | 0.1573 |
| 5th cerebellar lobule | 2.3 | 2.5 | < | 4.8 | 2.3 | 0.015 | 0.14453 |
| basal amygdala | 5.7 | 0.8 | > | 4.9 | 0.8 | 0.016 | 0.14185 |
| anterior olfactory n. | 16.4 | 2.9 | > | 14.1 | 2.1 | 0.019 | 0.13233 |
| paraventricular thalamus | 1.2 | 0.3 | > | 1.0 | 0.3 | 0.022 | 0.12559 |
| rostral piriform ctx | 14.2 | 2.9 | > | 11.9 | 1.7 | 0.025 | 0.11863 |
| extended amygdala | 2.4 | 1.0 | > | 1.8 | 0.6 | 0.026 | 0.11651 |
| orbital ctx | 17.7 | 5.9 | > | 12.6 | 3.9 | 0.026 | 0.11634 |
| trigeminal complex medulla | 2.6 | 1.8 | < | 4.1 | 1.7 | 0.035 | 0.10135 |
| primary motor ctx | 13.3 | 2.6 | > | 11.4 | 1.9 | 0.035 | 0.10126 |
| vestibular n. | 2.0 | 0.8 | > | 1.5 | 0.6 | 0.037 | 0.09922 |
| diagonal band of Broca | 0.4 | 0.3 | < | 0.6 | 0.3 | 0.038 | 0.09732 |
| red n. | 0.5 | 0.2 | < | 0.7 | 0.2 | 0.047 | 0.08738 |
| primary somatosensory ctx | 26.0 | 3.0 | > | 24.1 | 3.3 | 0.047 | 0.08697 |
| habenula n. | 0.5 | 0.2 | < | 0.7 | 0.2 | 0.049 | 0.08528 |

**Table 3.** The list of brain regions that were significantly different in volume between OXT males and Control males (FDR p=0.06). Males were most affected by OXT a birth showing 35/116 brain regions with significant differences from vehicle controls. As in the case of the females exposed to OXT at birth, the majority of the affected brain regions in OXT males were smaller than vehicle controls. The regions most sensitive were many of the same for females e.g. olfactory system, limbic cortex, basal ganglia, striatum, amygdala, and hypothalamus. The brain regions that were significantly larger in volume with OXT exposure were in the cerebellum (5th, 7th,8th, 9th lobules) and brainstem (gigantocellularis, trigeminal complex, cuneate nuc., medullary reticular nuc. pontine nuc.).

| Male Vehicle vs Male Oxytocin - Brain Volumes (mm3) |  |  |  |  |  |  |  |
| --- | --- | --- | --- | --- | --- | --- | --- |
| Brain Area | Male Veh |  |  | Male OT |  | P-val | ω Sq |
|  | Ave | SD |  | Ave | SD |  |  |
| frontal association ctx | 13.4 | 1.9 | > | 9.7 | 2.8 | 0.000 | 0.32943 |
| agranular insular ctx | 13.6 | 3.8 | > | 9.7 | 2.5 | 0.002 | 0.22789 |
| claustrum | 2.4 | 1.0 | > | 1.5 | 0.6 | 0.003 | 0.21273 |
| rostral piriform ctx | 13.2 | 2.4 | > | 10.5 | 2.3 | 0.003 | 0.20768 |
| anterior olfactory n. | 14.9 | 1.8 | > | 12.2 | 2.8 | 0.004 | 0.1931 |
| 5th cerebellar lobule | 2.7 | 2.8 | < | 5.4 | 2.1 | 0.005 | 0.19098 |
| endopiriform n. | 4.6 | 1.4 | > | 3.5 | 0.8 | 0.006 | 0.17671 |
| primary somatosensory ctx | 25.4 | 3.3 | > | 22.1 | 3.2 | 0.006 | 0.17439 |
| 7th cerebellar lobule | 0.3 | 0.9 | < | 2.3 | 2.8 | 0.007 | 0.17291 |
| accumbens core | 2.3 | 0.7 | > | 1.7 | 0.5 | 0.007 | 0.16991 |
| granular cell layer olfactory bulb | 11.4 | 3.5 | > | 9.0 | 1.2 | 0.008 | 0.16111 |
| orbital ctx | 15.7 | 6.0 | > | 11.0 | 2.8 | 0.011 | 0.14783 |
| globus pallidus | 3.5 | 0.6 | > | 2.8 | 0.6 | 0.012 | 0.14374 |
| caudate putamen (striatum) | 25.0 | 4.6 | > | 21.3 | 3.7 | 0.012 | 0.14359 |
| glomerular layer olfactory bulb | 18.9 | 4.9 | > | 15.2 | 2.9 | 0.013 | 0.13943 |
| cortical amygdala | 3.5 | 0.5 | > | 3.0 | 0.6 | 0.016 | 0.12943 |
| retrosplenial ctx | 7.6 | 1.1 | > | 6.7 | 1.0 | 0.016 | 0.12923 |
| 9th cerebellar lobule | 0.1 | 0.4 | < | 1.4 | 1.9 | 0.018 | 0.12523 |
| cuneate n. | 0.0 | 0.1 | < | 0.4 | 0.5 | 0.018 | 0.12523 |
| trigeminal complex | 3.1 | 2.6 | < | 4.5 | 1.5 | 0.019 | 0.12132 |
| secondary somatosensory ctx | 6.2 | 1.0 | > | 5.2 | 1.0 | 0.019 | 0.1213 |
| visual 2 ctx | 9.0 | 1.0 | > | 8.1 | 1.4 | 0.020 | 0.11941 |
| zona incerta | 1.6 | 0.2 | < | 1.8 | 0.3 | 0.022 | 0.11573 |
| dentate gyrus | 8.1 | 0.4 | > | 7.5 | 1.1 | 0.022 | 0.11556 |
| basal amygdala | 5.2 | 0.7 | > | 4.5 | 0.9 | 0.025 | 0.10986 |
| anterior cingulate ctx | 4.2 | 0.9 | > | 3.5 | 0.8 | 0.026 | 0.10789 |
| caudal piriform ctx | 9.3 | 1.1 | > | 8.3 | 1.2 | 0.027 | 0.10603 |
| anterior thalamus | 2.0 | 0.8 | > | 1.3 | 0.4 | 0.029 | 0.10239 |
| gigantocellular reticular n. | 4.2 | 2.4 | < | 5.8 | 1.6 | 0.031 | 0.09872 |
| central amygdaloid n. | 2.3 | 0.3 | > | 2.0 | 0.5 | 0.032 | 0.09704 |
| anterior hypothalamic area | 3.9 | 1.3 | > | 3.1 | 1.0 | 0.034 | 0.09515 |
| medullary reticular n. | 0.7 | 2.5 | < | 2.7 | 3.4 | 0.038 | 0.08917 |
| 8th cerebellar lobule | 0.3 | 1.1 | < | 1.6 | 1.9 | 0.039 | 0.08829 |
| pontine n. | 0.5 | 0.2 | < | 0.6 | 0.2 | 0.047 | 0.07992 |
| infralimbic ctx | 1.0 | 0.3 | > | 0.7 | 0.5 | 0.047 | 0.0797 |

**Table 4.**
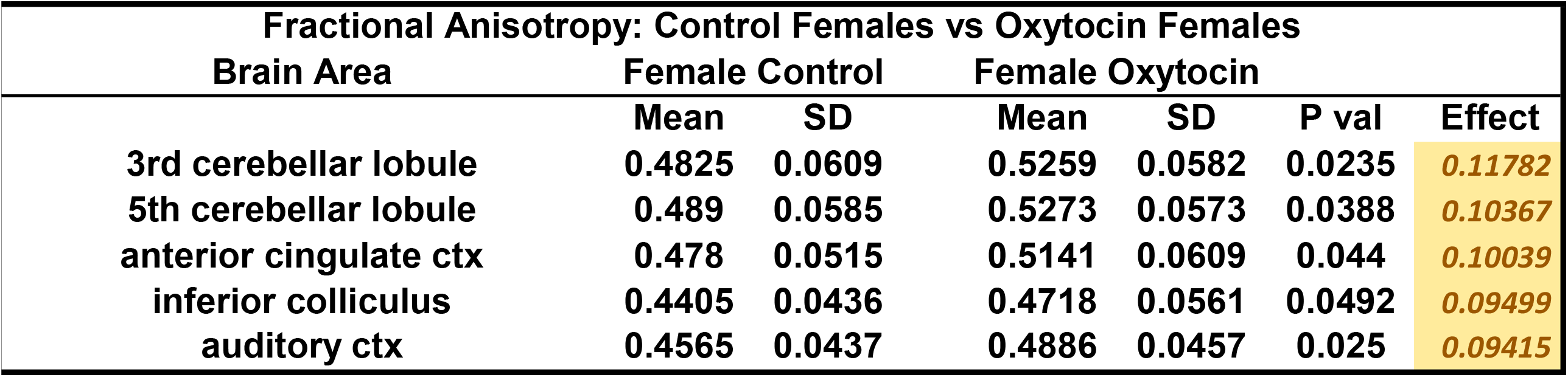
The list of brain regions that were significantly different in fractional anisotropy between OXT females and Control females.

| Fractional Anisotropy: Control Females vs Oxytocin Females |  |  |  |  |  |  |
| --- | --- | --- | --- | --- | --- | --- |
| Brain Area | Female Control |  | Female Oxytocin |  | P val | Effect |
|  | Mean | SD | Mean | SD |  |  |
| 3rd cerebellar lobule | 0.4825 | 0.0609 | 0.5259 | 0.0582 | 0.0235 | 0.11782 |
| 5th cerebellar lobule | 0.489 | 0.0585 | 0.5273 | 0.0573 | 0.0388 | 0.10367 |
| anterior cingulate ctx | 0.478 | 0.0515 | 0.5141 | 0.0609 | 0.044 | 0.10039 |
| inferior colliculus | 0.4405 | 0.0436 | 0.4718 | 0.0561 | 0.0492 | 0.09499 |
| auditory ctx | 0.4565 | 0.0437 | 0.4886 | 0.0457 | 0.025 | 0.09415 |

**Figure 1.**
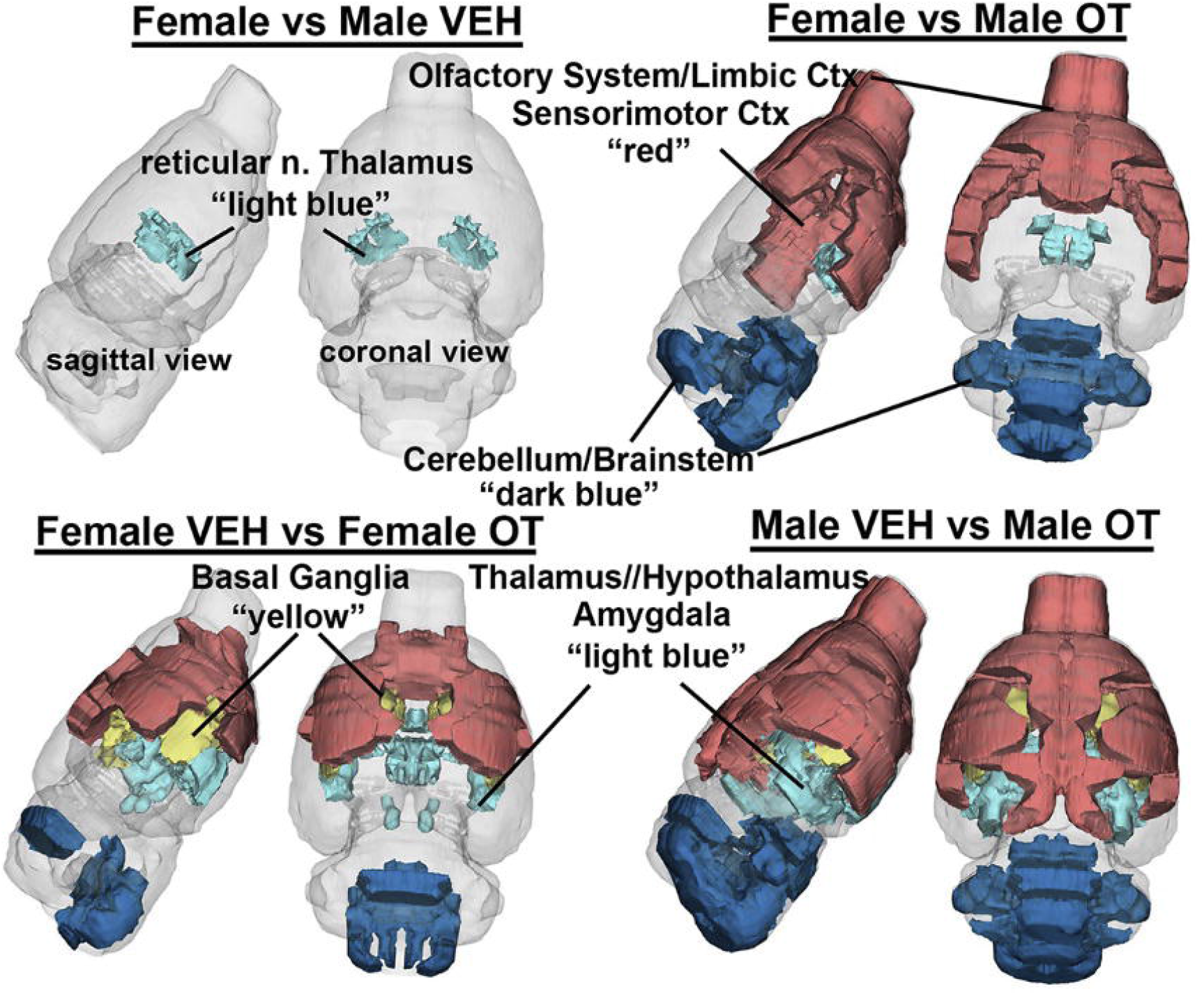
A 3D color coded reconstructions summarizing the significantly different brain areas with volumetric changes for each experimental condition. Details of these differences can be found in tables 1-3.

**Figure 2.**
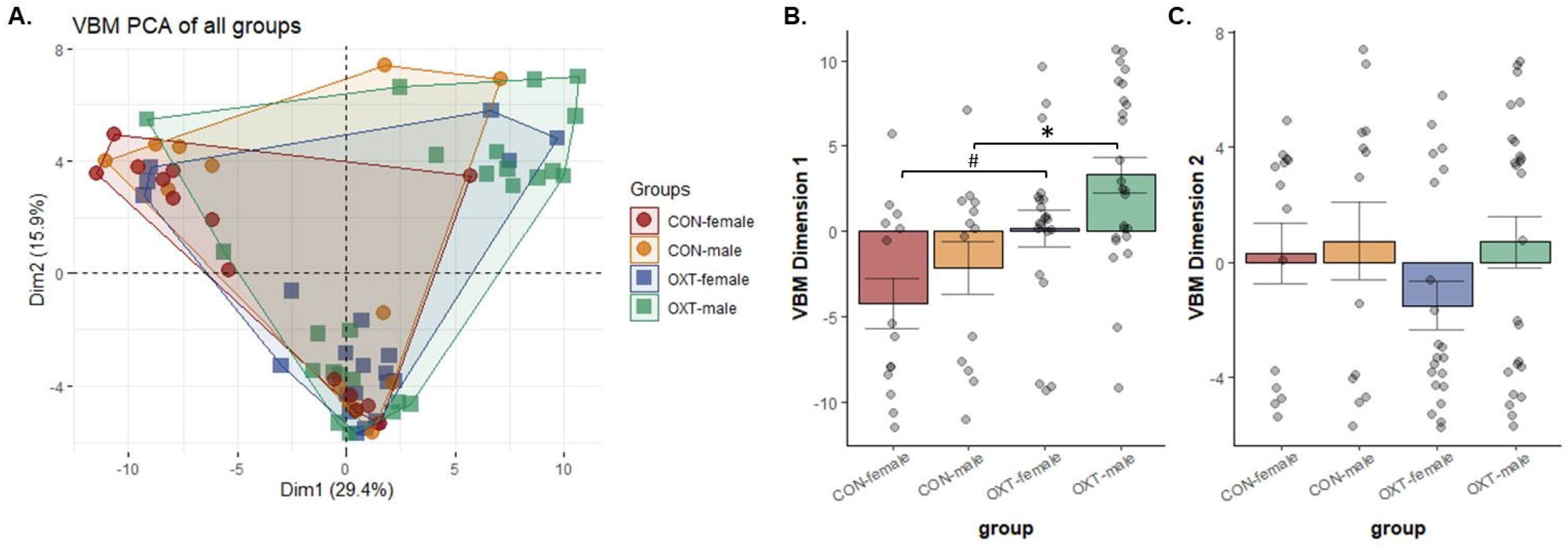
(A) Voxel-based morphometry (VBM) measures from 111 brain regions were loaded into a principal component analysis. The overall explanatory value of dimensions 1 and 2 was 29.4% and 15.9%, respectively. (B) Both male sex and OXT treatment lead to greater values in dimension 1 (p < 0.029 for both comparisons). Post-hoc analyses revealed OXT males had significantly greater dimension 1 scores than Control males (* p = 0.017), while OXT females tended to be greater than Control females (# p = 0.079). (C) There were no effects in dimension 2.

We observed a broad, albeit subtle pattern of effects in OXT-exposed males in DWI measures. In terms of FA, Control animals showed small but widespread sex differences, with females having greater FA than males across brain regions, a pattern not present in OXT animals due to increased FA among OXT-exposed males (Figure 3A, Tables 2 and 3). Indeed, there were no differences in either FA or ADC between OXT males and OXT females, whereas there were 62 and 48 such regions amongst Control animals. When FA data from all 111 brain regions were loaded into a PCA, there were no effects of either sex or treatment detected (Figure 3A). In terms of ADC, Control males had greater ADC values than OXT males across 84 brain regions (Table 5). While we observed widespread ADC differences between Control males and females, we observed no sex differences in ADC in the OXT-exposed condition. The PCA for ADC revealed a main effect of OXT exposure (F(1,65) = 5.80, p = 0.019) and a trend toward an interaction between sex and treatment (p = 0.087). Post-hoc analysis revealed that Control males having greater dimension 1 values than both OXT males (p = 0.026) and OXT females (p = 0.042). Thus, across the brain, OXT males’ ADC values more closely resembled Control females and OXT females than they did Control males.

**Figure 3.**
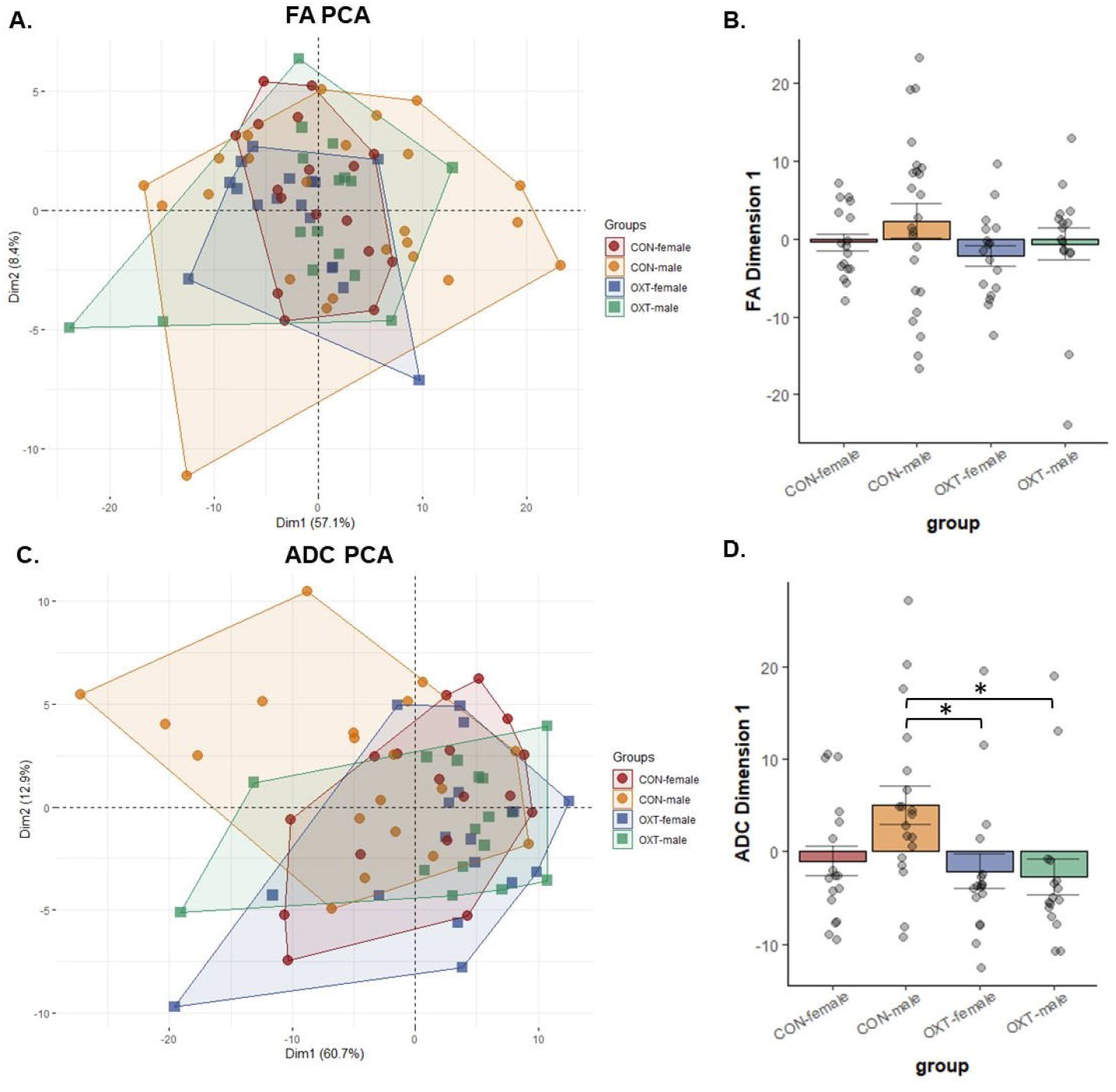
Diffusion-weighted imaging (DWI) measures for fractional anisotropy (FA, panels A and B) and apparent diffusion coefficient (ADC, panels C and D) from 111 brain regions were loaded into a principal component analysis. (B) There were no significant differences in FA. (D) Control males had greater dimension 1 scores than OXT males (* p = 0.026) and OXT females (* p = 0.042). in terms of ADC.

**Table 5.**
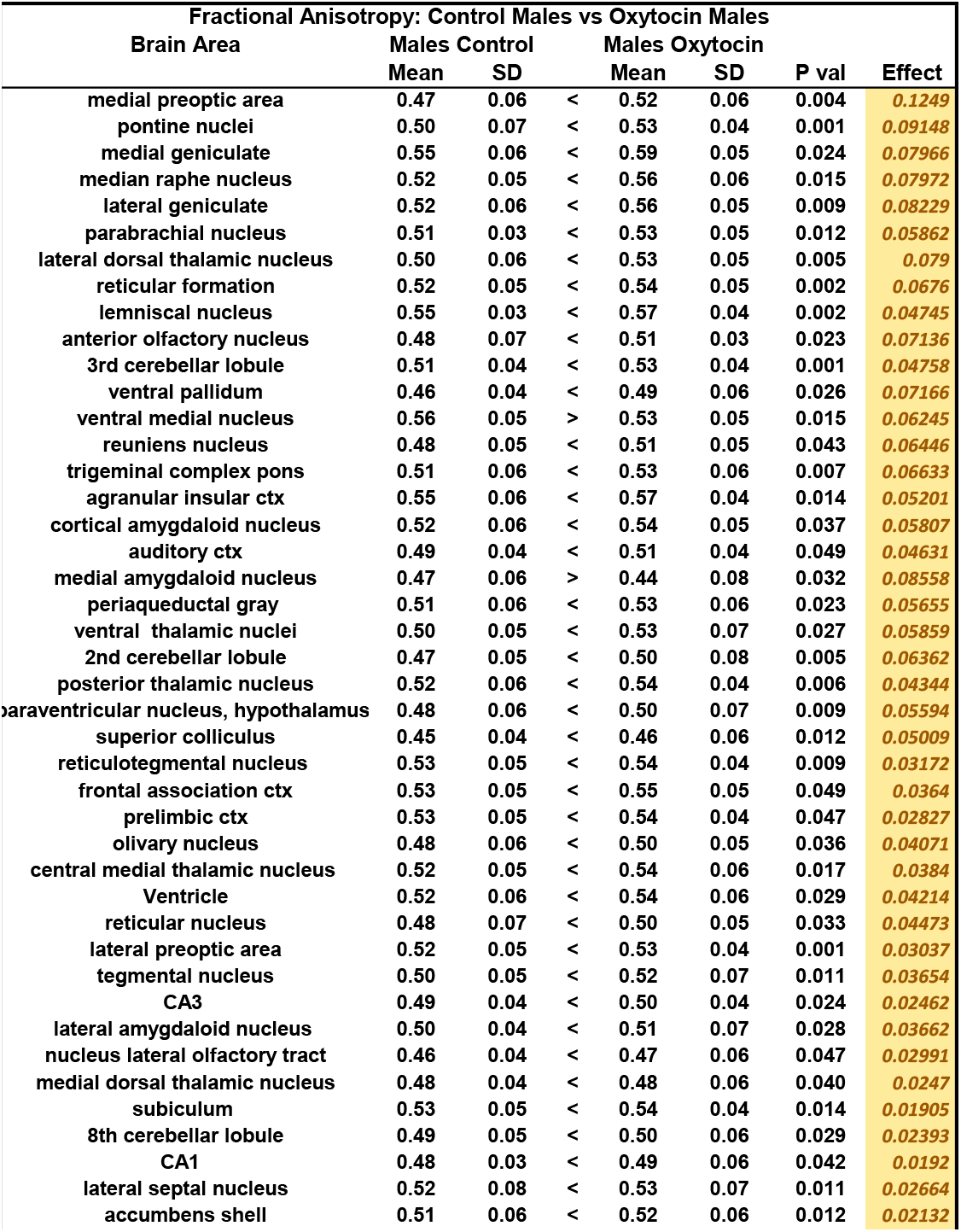
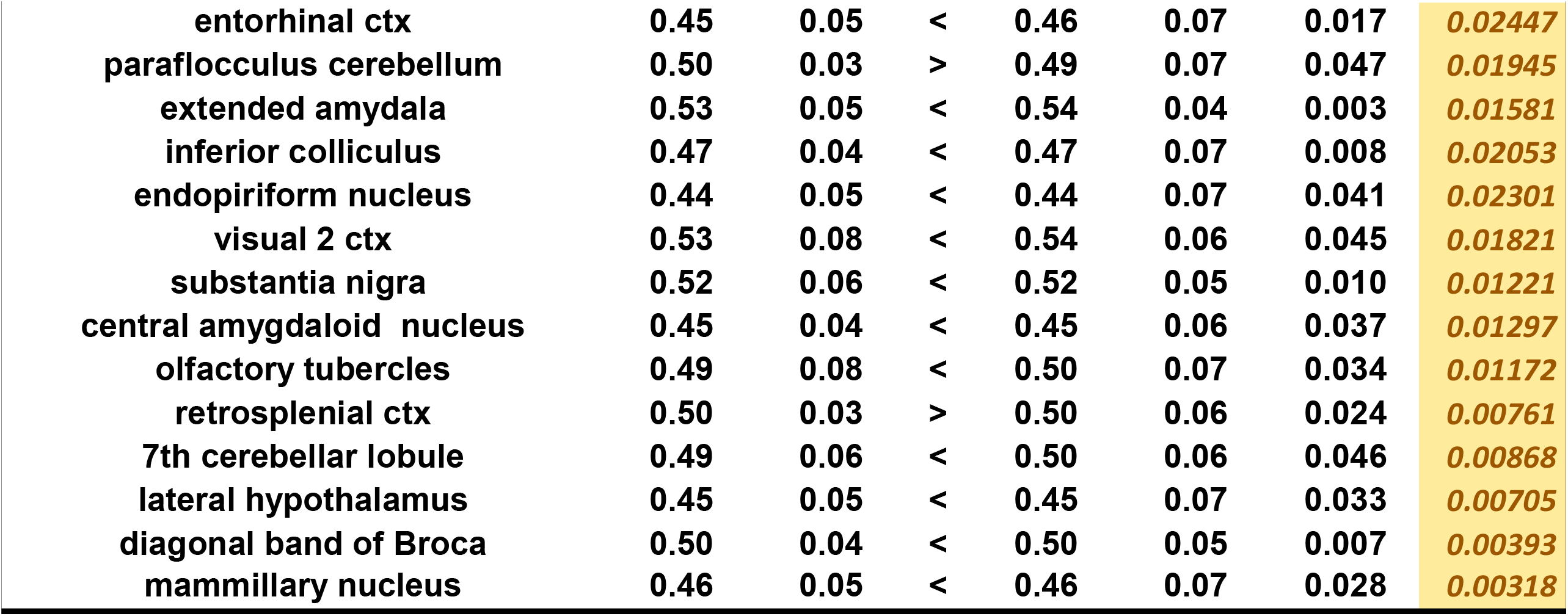
The list of brain regions that were significantly different in fractional anisotropy (FA) between OXT males and Control males. While widespread, differences were generally small in effect, with OXT males generally showing greater FA scores.

| Fractional Anisotropy: Control Males vs Oxytocin Males |  |  |  |  |  |  |  |
| --- | --- | --- | --- | --- | --- | --- | --- |
| Brain Area | Males Control |  |  | Males Oxytocin |  | P val | Effect |
|  | Mean | SD |  | Mean | SD |  |  |
| medial preoptic area | 0.47 | 0.06 | < | 0.52 | 0.06 | 0.004 | 0.1249 |
| pontine nuclei | 0.50 | 0.07 | < | 0.53 | 0.04 | 0.001 | 0.09148 |
| medial geniculate | 0.55 | 0.06 | < | 0.59 | 0.05 | 0.024 | 0.07966 |
| median raphe nucleus | 0.52 | 0.05 | < | 0.56 | 0.06 | 0.015 | 0.07972 |
| lateral geniculate | 0.52 | 0.06 | < | 0.56 | 0.05 | 0.009 | 0.08229 |
| parabrachial nucleus | 0.51 | 0.03 | < | 0.53 | 0.05 | 0.012 | 0.05862 |
| lateral dorsal thalamic nucleus | 0.50 | 0.06 | < | 0.53 | 0.05 | 0.005 | 0.079 |
| reticular formation | 0.52 | 0.05 | < | 0.54 | 0.05 | 0.002 | 0.0676 |
| lemniscal nucleus | 0.55 | 0.03 | < | 0.57 | 0.04 | 0.002 | 0.04745 |
| anterior olfactory nucleus | 0.48 | 0.07 | < | 0.51 | 0.03 | 0.023 | 0.07136 |
| 3rd cerebellar lobule | 0.51 | 0.04 | < | 0.53 | 0.04 | 0.001 | 0.04758 |
| ventral pallidum | 0.46 | 0.04 | < | 0.49 | 0.06 | 0.026 | 0.07166 |
| ventral medial nucleus | 0.56 | 0.05 | > | 0.53 | 0.05 | 0.015 | 0.06245 |
| reuniens nucleus | 0.48 | 0.05 | < | 0.51 | 0.05 | 0.043 | 0.06446 |
| trigeminal complex pons | 0.51 | 0.06 | < | 0.53 | 0.06 | 0.007 | 0.06633 |
| agranular insular ctx | 0.55 | 0.06 | < | 0.57 | 0.04 | 0.014 | 0.05201 |
| cortical amygdaloid nucleus | 0.52 | 0.06 | < | 0.54 | 0.05 | 0.037 | 0.05807 |
| auditory ctx | 0.49 | 0.04 | < | 0.51 | 0.04 | 0.049 | 0.04631 |
| medial amygdaloid nucleus | 0.47 | 0.06 | > | 0.44 | 0.08 | 0.032 | 0.08558 |
| periaqueductal gray | 0.51 | 0.06 | < | 0.53 | 0.06 | 0.023 | 0.05655 |
| ventral thalamic nuclei | 0.50 | 0.05 | < | 0.53 | 0.07 | 0.027 | 0.05859 |
| 2nd cerebellar lobule | 0.47 | 0.05 | < | 0.50 | 0.08 | 0.005 | 0.06362 |
| posterior thalamic nucleus | 0.52 | 0.06 | < | 0.54 | 0.04 | 0.006 | 0.04344 |
| paraventricular nucleus, hypothalamus | 0.48 | 0.06 | < | 0.50 | 0.07 | 0.009 | 0.05594 |
| superior colliculus | 0.45 | 0.04 | < | 0.46 | 0.06 | 0.012 | 0.05009 |
| reticulotegmental nucleus | 0.53 | 0.05 | < | 0.54 | 0.04 | 0.009 | 0.03172 |
| frontal association ctx | 0.53 | 0.05 | < | 0.55 | 0.05 | 0.049 | 0.0364 |
| prelimbic ctx | 0.53 | 0.05 | < | 0.54 | 0.04 | 0.047 | 0.02827 |
| olivary nucleus | 0.48 | 0.06 | < | 0.50 | 0.05 | 0.036 | 0.04071 |
| central medial thalamic nucleus | 0.52 | 0.05 | < | 0.54 | 0.06 | 0.017 | 0.0384 |
| Ventricle | 0.52 | 0.06 | < | 0.54 | 0.06 | 0.029 | 0.04214 |
| reticular nucleus | 0.48 | 0.07 | < | 0.50 | 0.05 | 0.033 | 0.04473 |
| lateral preoptic area | 0.52 | 0.05 | < | 0.53 | 0.04 | 0.001 | 0.03037 |
| tegmental nucleus | 0.50 | 0.05 | < | 0.52 | 0.07 | 0.011 | 0.03654 |
| CA3 | 0.49 | 0.04 | < | 0.50 | 0.04 | 0.024 | 0.02462 |
| lateral amygdaloid nucleus | 0.50 | 0.04 | < | 0.51 | 0.07 | 0.028 | 0.03662 |
| nucleus lateral olfactory tract | 0.46 | 0.04 | < | 0.47 | 0.06 | 0.047 | 0.02991 |
| medial dorsal thalamic nucleus | 0.48 | 0.04 | < | 0.48 | 0.06 | 0.040 | 0.0247 |
| subiculum | 0.53 | 0.05 | < | 0.54 | 0.04 | 0.014 | 0.01905 |
| 8th cerebellar lobule | 0.49 | 0.05 | < | 0.50 | 0.06 | 0.029 | 0.02393 |
| CA1 | 0.48 | 0.03 | < | 0.49 | 0.06 | 0.042 | 0.0192 |
| lateral septal nucleus | 0.52 | 0.08 | < | 0.53 | 0.07 | 0.011 | 0.02664 |
| accumbens shell | 0.51 | 0.06 | < | 0.52 | 0.06 | 0.012 | 0.02132 |
| entorhinal ctx | 0.45 | 0.05 | < | 0.46 | 0.07 | 0.017 | 0.02447 |
| paraflocculus cerebellum | 0.50 | 0.03 | > | 0.49 | 0.07 | 0.047 | 0.01945 |
| extended amygdala | 0.53 | 0.05 | < | 0.54 | 0.04 | 0.003 | 0.01581 |
| inferior colliculus | 0.47 | 0.04 | < | 0.47 | 0.07 | 0.008 | 0.02053 |
| endopiriform nucleus | 0.44 | 0.05 | < | 0.44 | 0.07 | 0.041 | 0.02301 |
| visual 2 ctx | 0.53 | 0.08 | < | 0.54 | 0.06 | 0.045 | 0.01821 |
| substantia nigra | 0.52 | 0.06 | < | 0.52 | 0.05 | 0.010 | 0.01221 |
| central amygdaloid nucleus | 0.45 | 0.04 | < | 0.45 | 0.06 | 0.037 | 0.01297 |
| olfactory tubercles | 0.49 | 0.08 | < | 0.50 | 0.07 | 0.034 | 0.01172 |
| retrosplenial ctx | 0.50 | 0.03 | > | 0.50 | 0.06 | 0.024 | 0.00761 |
| 7th cerebellar lobule | 0.49 | 0.06 | < | 0.50 | 0.06 | 0.046 | 0.00868 |
| lateral hypothalamus | 0.45 | 0.05 | < | 0.45 | 0.07 | 0.033 | 0.00705 |
| diagonal band of Broca | 0.50 | 0.04 | < | 0.50 | 0.05 | 0.007 | 0.00393 |
| mammillary nucleus | 0.46 | 0.05 | < | 0.46 | 0.07 | 0.028 | 0.00318 |

**Table 6.**
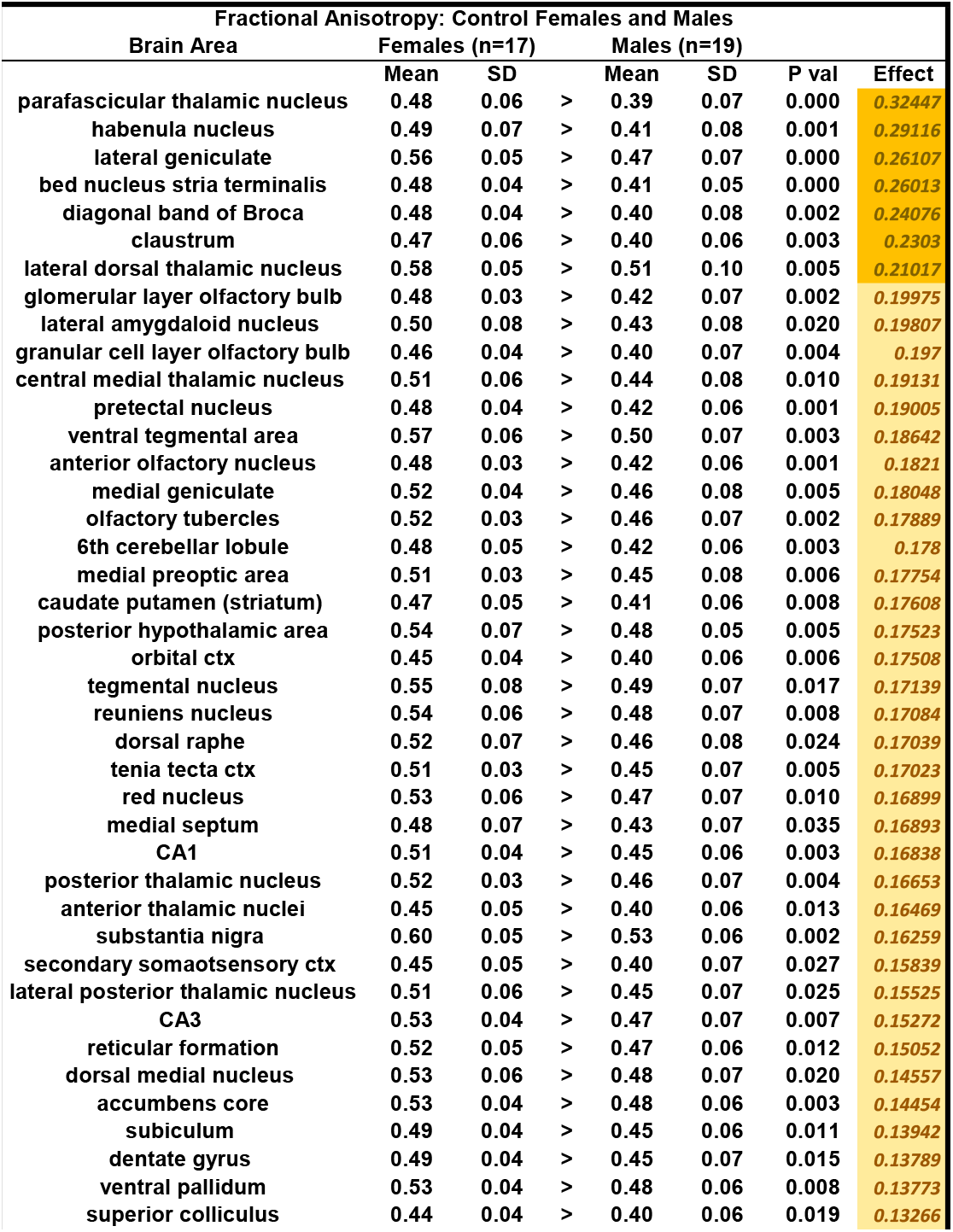
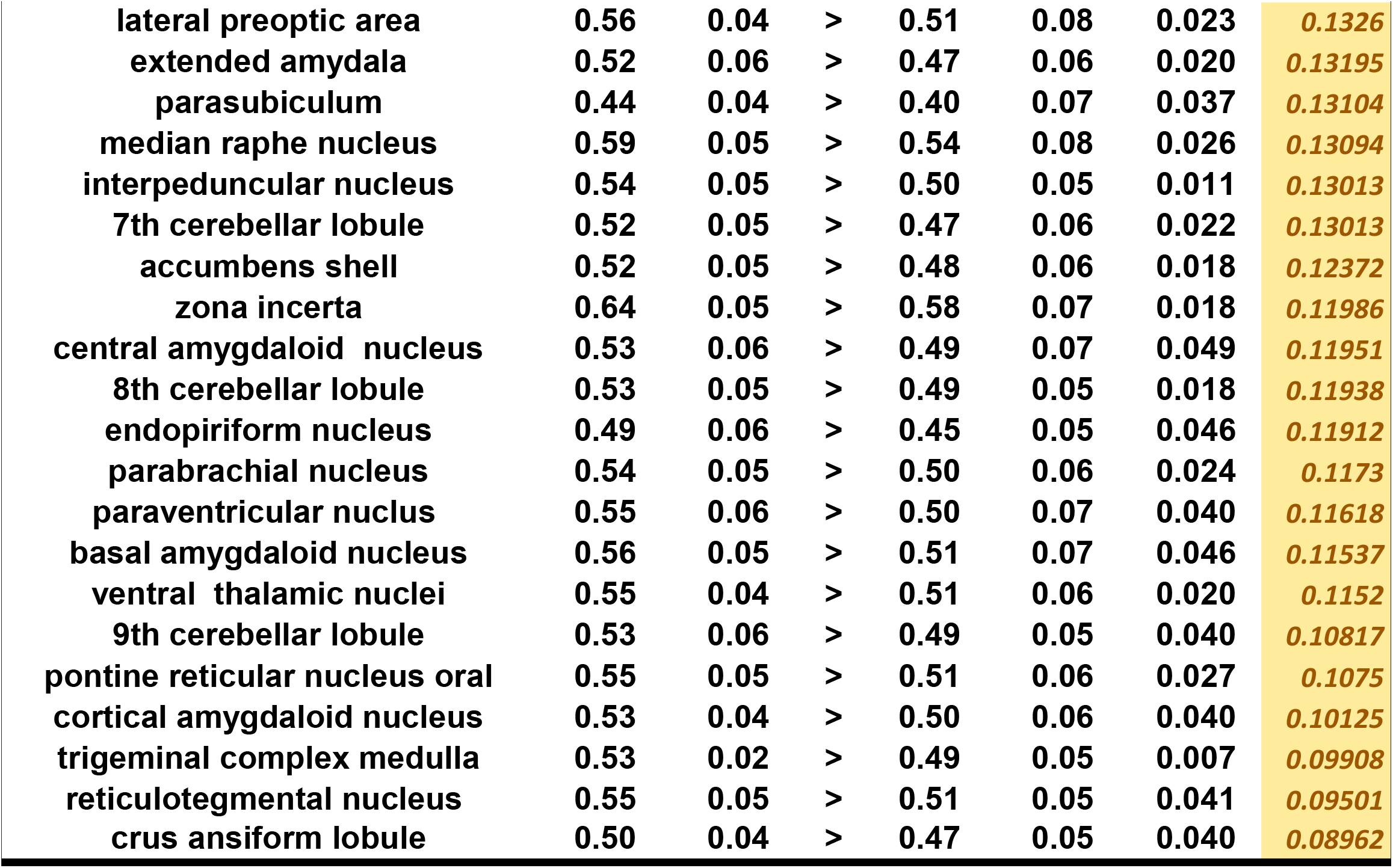
The list of brain regions that were significantly different in fractional anisotropy (FA) between Control females and Control males. While widespread, differences outside of the thalamus were generally small in effect, with females generally showing greater FA scores.

Fractional Anisotropy: Control Females and Males
| Brain Area | Females (n=17) |  |  | Males (n=19) |  | P val | Effect |
| --- | --- | --- | --- | --- | --- | --- | --- |
|  | Mean | SD |  | Mean | SD |  |  |
| parafascicular thalamic nucleus | 0.48 | 0.06 | > | 0.39 | 0.07 | 0.000 | 0.32447 |
| habenula nucleus | 0.49 | 0.07 | > | 0.41 | 0.08 | 0.001 | 0.29116 |
| lateral geniculate | 0.56 | 0.05 | > | 0.47 | 0.07 | 0.000 | 0.26107 |
| bed nucleus stria terminalis | 0.48 | 0.04 | > | 0.41 | 0.05 | 0.000 | 0.26013 |
| diagonal band of Broca | 0.48 | 0.04 | > | 0.40 | 0.08 | 0.002 | 0.24076 |
| claustrum | 0.47 | 0.06 | > | 0.40 | 0.06 | 0.003 | 0.2303 |
| lateral dorsal thalamic nucleus | 0.58 | 0.05 | > | 0.51 | 0.10 | 0.005 | 0.21017 |
| glomerular layer olfactory bulb | 0.48 | 0.03 | > | 0.42 | 0.07 | 0.002 | 0.19975 |
| lateral amygdaloid nucleus | 0.50 | 0.08 | > | 0.43 | 0.08 | 0.020 | 0.19807 |
| granular cell layer olfactory bulb | 0.46 | 0.04 | > | 0.40 | 0.07 | 0.004 | 0.197 |
| central medial thalamic nucleus | 0.51 | 0.06 | > | 0.44 | 0.08 | 0.010 | 0.19131 |
| pretectal nucleus | 0.48 | 0.04 | > | 0.42 | 0.06 | 0.001 | 0.19005 |
| ventral tegmental area | 0.57 | 0.06 | > | 0.50 | 0.07 | 0.003 | 0.18642 |
| anterior olfactory nucleus | 0.48 | 0.03 | > | 0.42 | 0.06 | 0.001 | 0.1821 |
| medial geniculate | 0.52 | 0.04 | > | 0.46 | 0.08 | 0.005 | 0.18048 |
| olfactory tubercles | 0.52 | 0.03 | > | 0.46 | 0.07 | 0.002 | 0.17889 |
| 6th cerebellar lobule | 0.48 | 0.05 | > | 0.42 | 0.06 | 0.003 | 0.178 |
| medial preoptic area | 0.51 | 0.03 | > | 0.45 | 0.08 | 0.006 | 0.17754 |
| caudate putamen (striatum) | 0.47 | 0.05 | > | 0.41 | 0.06 | 0.008 | 0.17608 |
| posterior hypothalamic area | 0.54 | 0.07 | > | 0.48 | 0.05 | 0.005 | 0.17523 |
| orbital ctx | 0.45 | 0.04 | > | 0.40 | 0.06 | 0.006 | 0.17508 |
| tegmental nucleus | 0.55 | 0.08 | > | 0.49 | 0.07 | 0.017 | 0.17139 |
| reuniens nucleus | 0.54 | 0.06 | > | 0.48 | 0.07 | 0.008 | 0.17084 |
| dorsal raphe | 0.52 | 0.07 | > | 0.46 | 0.08 | 0.024 | 0.17039 |
| tenia tecta ctx | 0.51 | 0.03 | > | 0.45 | 0.07 | 0.005 | 0.17023 |
| red nucleus | 0.53 | 0.06 | > | 0.47 | 0.07 | 0.010 | 0.16899 |
| medial septum | 0.48 | 0.07 | > | 0.43 | 0.07 | 0.035 | 0.16893 |
| CA1 | 0.51 | 0.04 | > | 0.45 | 0.06 | 0.003 | 0.16838 |
| posterior thalamic nucleus | 0.52 | 0.03 | > | 0.46 | 0.07 | 0.004 | 0.16653 |
| anterior thalamic nuclei | 0.45 | 0.05 | > | 0.40 | 0.06 | 0.013 | 0.16469 |
| substantia nigra | 0.60 | 0.05 | > | 0.53 | 0.06 | 0.002 | 0.16259 |
| secondary somatosensory ctx | 0.45 | 0.05 | > | 0.40 | 0.07 | 0.027 | 0.15839 |
| lateral posterior thalamic nucleus | 0.51 | 0.06 | > | 0.45 | 0.07 | 0.025 | 0.15525 |
| CA3 | 0.53 | 0.04 | > | 0.47 | 0.07 | 0.007 | 0.15272 |
| reticular formation | 0.52 | 0.05 | > | 0.47 | 0.06 | 0.012 | 0.15052 |
| dorsal medial nucleus | 0.53 | 0.06 | > | 0.48 | 0.07 | 0.020 | 0.14557 |
| accumbens core | 0.53 | 0.04 | > | 0.48 | 0.06 | 0.003 | 0.14454 |
| subiculum | 0.49 | 0.04 | > | 0.45 | 0.06 | 0.011 | 0.13942 |
| dentate gyrus | 0.49 | 0.04 | > | 0.45 | 0.07 | 0.015 | 0.13789 |
| ventral pallidum | 0.53 | 0.04 | > | 0.48 | 0.06 | 0.008 | 0.13773 |
| superior colliculus | 0.44 | 0.04 | > | 0.40 | 0.06 | 0.019 | 0.13266 |
| lateral preoptic area | 0.56 | 0.04 | > | 0.51 | 0.08 | 0.023 | 0.1326 |
| extended amygdala | 0.52 | 0.06 | > | 0.47 | 0.06 | 0.020 | 0.13195 |
| parasubiculum | 0.44 | 0.04 | > | 0.40 | 0.07 | 0.037 | 0.13104 |
| median raphe nucleus | 0.59 | 0.05 | > | 0.54 | 0.08 | 0.026 | 0.13094 |
| interpeduncular nucleus | 0.54 | 0.05 | > | 0.50 | 0.05 | 0.011 | 0.13013 |
| 7th cerebellar lobule | 0.52 | 0.05 | > | 0.47 | 0.06 | 0.022 | 0.13013 |
| accumbens shell | 0.52 | 0.05 | > | 0.48 | 0.06 | 0.018 | 0.12372 |
| zona incerta | 0.64 | 0.05 | > | 0.58 | 0.07 | 0.018 | 0.11986 |
| central amygdaloid nucleus | 0.53 | 0.06 | > | 0.49 | 0.07 | 0.049 | 0.11951 |
| 8th cerebellar lobule | 0.53 | 0.05 | > | 0.49 | 0.05 | 0.018 | 0.11938 |
| endopiriform nucleus | 0.49 | 0.06 | > | 0.45 | 0.05 | 0.046 | 0.11912 |
| parabrachial nucleus | 0.54 | 0.05 | > | 0.50 | 0.06 | 0.024 | 0.1173 |
| paraventricular nucleus | 0.55 | 0.06 | > | 0.50 | 0.07 | 0.040 | 0.11618 |
| basal amygdaloid nucleus | 0.56 | 0.05 | > | 0.51 | 0.07 | 0.046 | 0.11537 |
| ventral thalamic nuclei | 0.55 | 0.04 | > | 0.51 | 0.06 | 0.020 | 0.1152 |
| 9th cerebellar lobule | 0.53 | 0.06 | > | 0.49 | 0.05 | 0.040 | 0.10817 |
| pontine reticular nucleus oral | 0.55 | 0.05 | > | 0.51 | 0.06 | 0.027 | 0.1075 |
| cortical amygdaloid nucleus | 0.53 | 0.04 | > | 0.50 | 0.06 | 0.040 | 0.10125 |
| trigeminal complex medulla | 0.53 | 0.02 | > | 0.49 | 0.05 | 0.007 | 0.09908 |
| reticulotegmental nucleus | 0.55 | 0.05 | > | 0.51 | 0.05 | 0.041 | 0.09501 |
| crus ansiform lobule | 0.50 | 0.04 | > | 0.47 | 0.05 | 0.040 | 0.08962 |

In the analyses of functional connectivity (Figures 4-8), OXT males stood out as having a widespread pattern of greater connectivity. The vast majority of connections across all groups arose from positive correlations. Male sex and OXT treatment both increased the proportion of significant connections for both positive and negative connections (chi-square p < 0.001 for both effects). Whereas Control females were found to have significant functional connectivity in 5% of all possible connections, OXT females had significant connectivity in 7.2% of connections. Whereas Control males had connectivity in 6.3% of connections, OXT males had significant connectivity in 12.5% of connections (Figure 4A). Similarly, male sex and OXT treatment at birth both increased the strength of connectivity among region-region pairs whose activity was significantly correlated, but only for positively correlated pairs (p < 0.001 for both effects, Figure 4B). Thus, OXT led to more regions significantly functionally connected and stronger correlations in such regions among males and to a lesser extent among females (Figures 4,5). In examining the pattern of results within correlation matrices, we observed a concentration of stronger connectivity among regions of the same cluster (Figure 4D and E).

**Figure 4.**
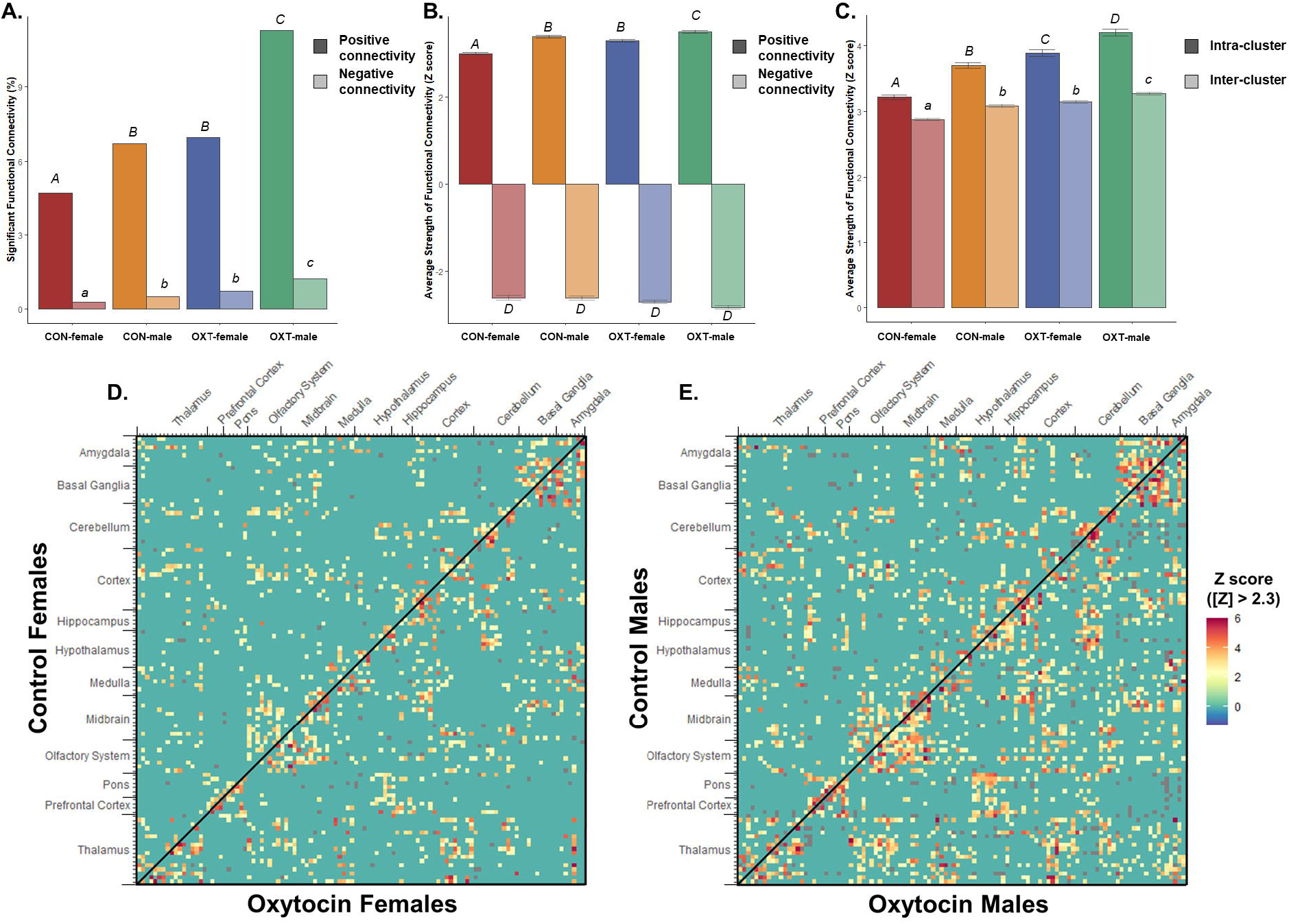
Resting-state functional connectivity from 111 brain regions. (A) Both OXT treatment and male sex increased functional connectivity, meaning OXT males had the greatest proportion of region-region pairs significantly functionally connected for both positive (dark fill) and negative (light fill) correlations. Significant group differences are indicated with different letters over top the bars. (B) The average strength of correlation among region-region pairs whose activity was significantly correlated. Both OXT treatment and male sex increased the strength of connectivity for positive connections (dark fill). There were no significant differences among negative connections (light fill). (C) Both OXT treatment and male sex increased the strength of connectivity among both intra- and inter-cluster, though intra-cluster connectivity was more sensitive to these effects. Significant group differences are indicated with different letters over top the bars. In panels (D) and (E), connectivity from males and females respectively, 111×111 cell matrices show the strength of connectivity for all possible pairs of brain regions. Reflected across the diagonal are opposing treatment conditions, with OXT on top and Control on bottom. Region-region pairs whose connectivity Z score was less than |2.3| were excluded.

**Figure 5.**
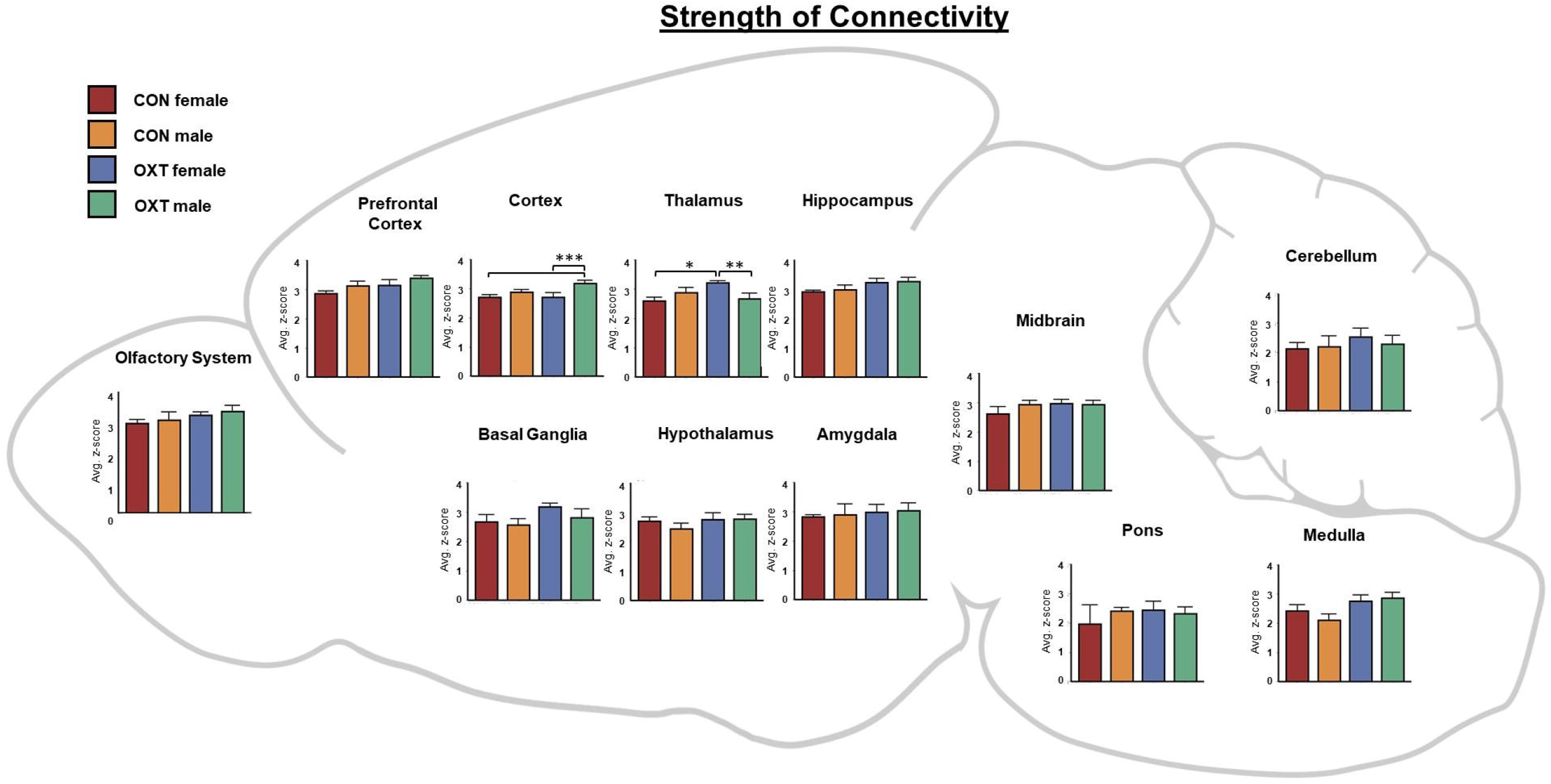
A map of the strength of connectivity (i.e. correlations’ z-scores) averaged over regional clusters by group. For example, the ‘Hippocampus’ cluster includes the: CA1, CA3, Dentate gyrus, Subiculum and Parasubiculum. * p < 0.05, ** p < 0.01, *** p < 0.001.

This led us to examine the strength of intra- vs. inter-cluster connectivity as a function of sex and birth treatment, which revealed that male sex and OXT treatment increased intra-cluster connectivity (e.g. central amygdala and medial amygdala) to a greater degree than inter-cluster connectivity (e.g. central amygdala and dentate gyrus, Figure 4C). As shown in Figure 5, when comparing the strength of connectivity at the level of regional clusters (e.g. all subregions of the amygdala), we observed a treatment effect in the medulla and olfactory system, however there were no significant post-hoc differences. There was also two treatment by sex interactions such that OXT males had weaker connectivity than OXT females across the thalamus (p < 0.002) and stronger connectivity than OXT females across the cortex (p < 0.001).

We next examined three indices from analyses based in graph theory: betweenness, closeness, and degree (see above for definitions of each). In terms of *betweenness*, male sex and OXT treatment both increased betweenness in the basal ganglia (Figure 6, p < 0.01 for both effects). In terms of *closeness*, we observed similar albeit more widespread effects, with male sex and OXT treatment and both increasing closeness across several regional clusters (Figure 7). Lastly, similar effects were found in terms of *degree*, with OXT males having greater degree across a wide swath of regional clusters (Figure 8).

**Figure 6.**
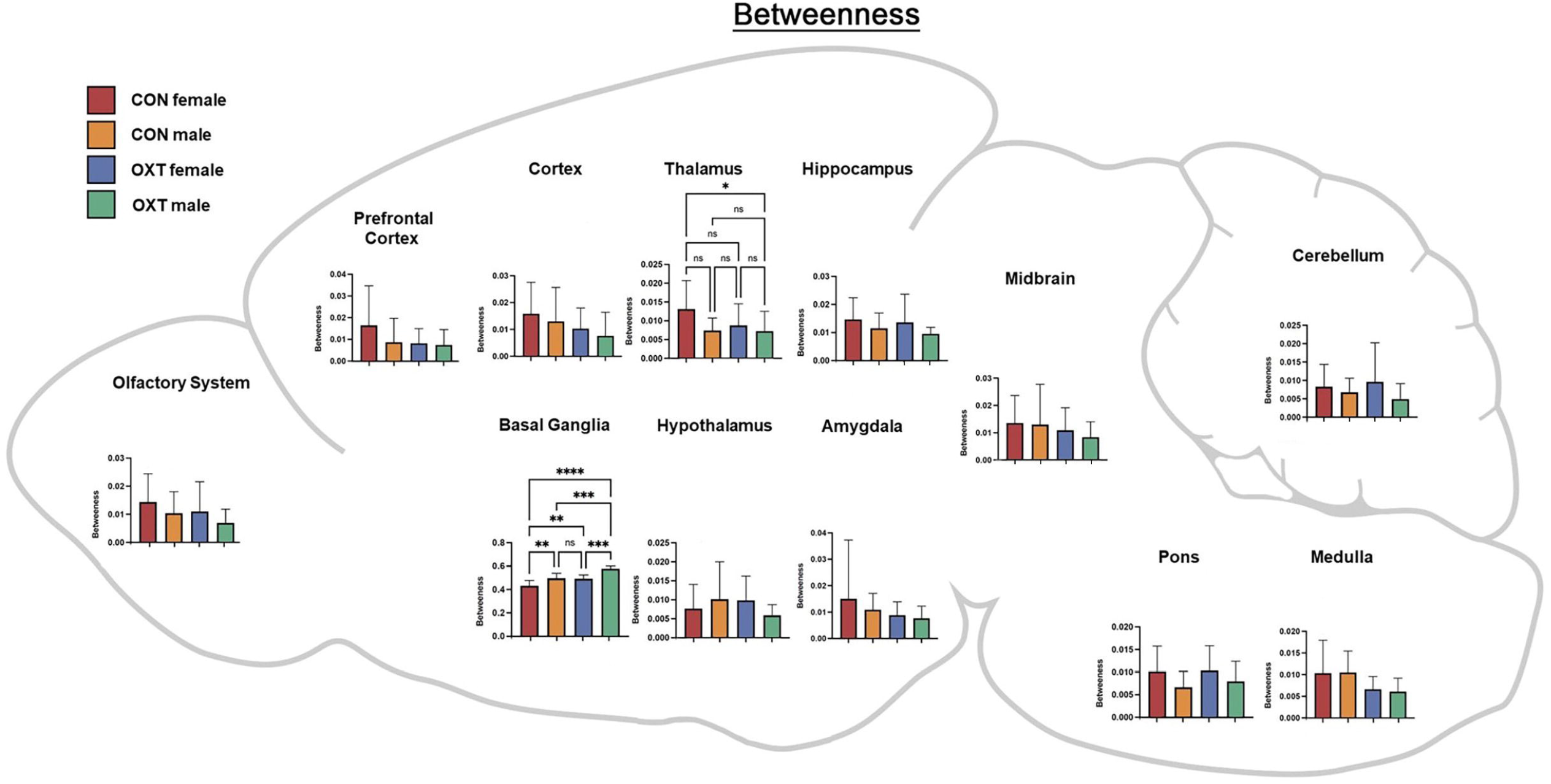
A map of betweenness averaged over regional clusters by group. For example, the ‘Hippocampus’ cluster includes the: CA1, CA3, Dentate gyrus, Subiculum and Parasubiculum. * p < 0.0332, ** p < 0.0021, *** p < 0.0002, **** p <0.0001.

**Figure 7.**
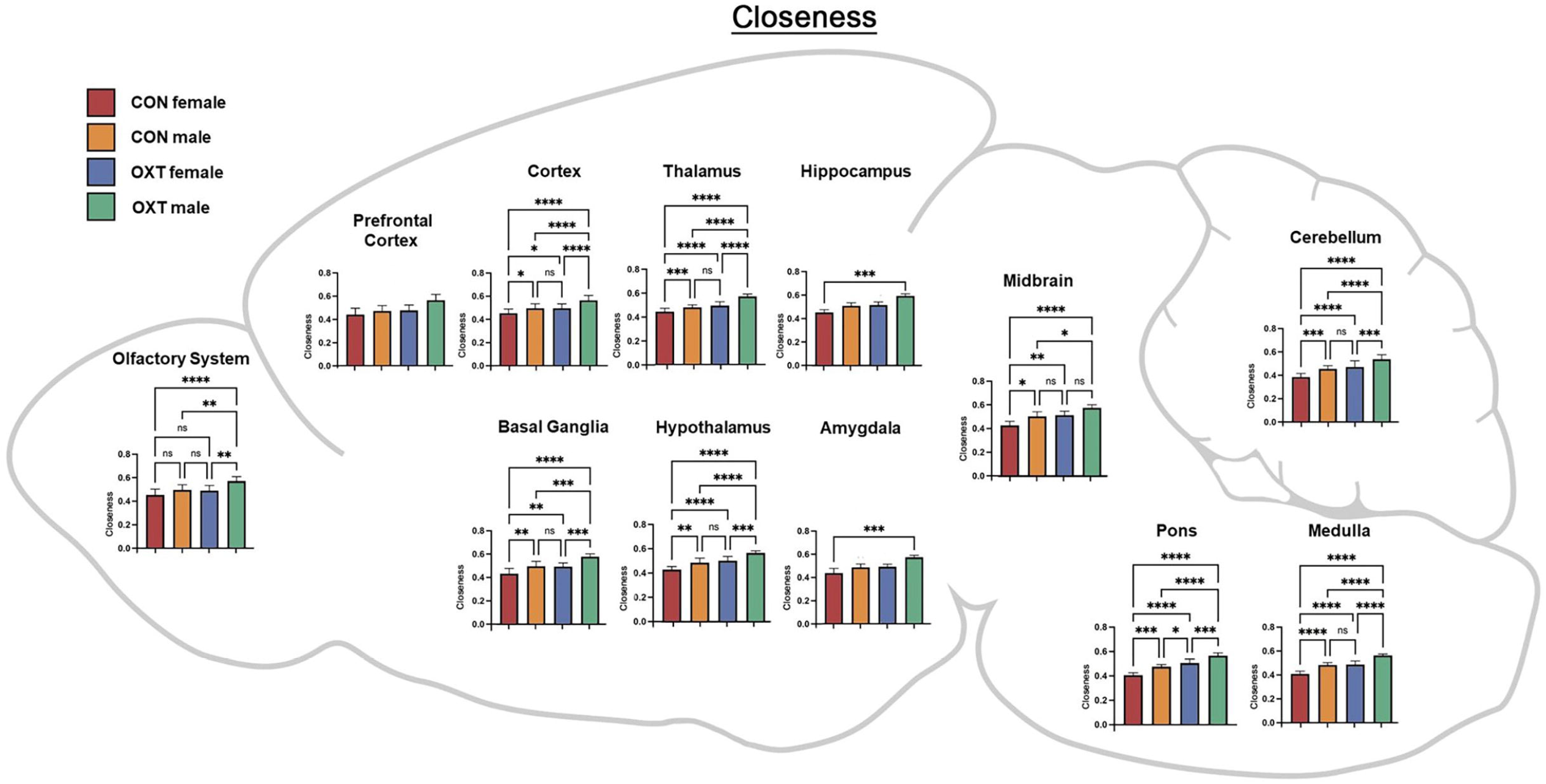
A map of closeness averaged over regional clusters by group. Both OXT treatment and male sex increased closeness throughout the brain, with effects most apparent in OXT males. * p < 0.0332, ** p < 0.0021, *** p < 0.0002, **** p <0.0001.

**Figure 8.**
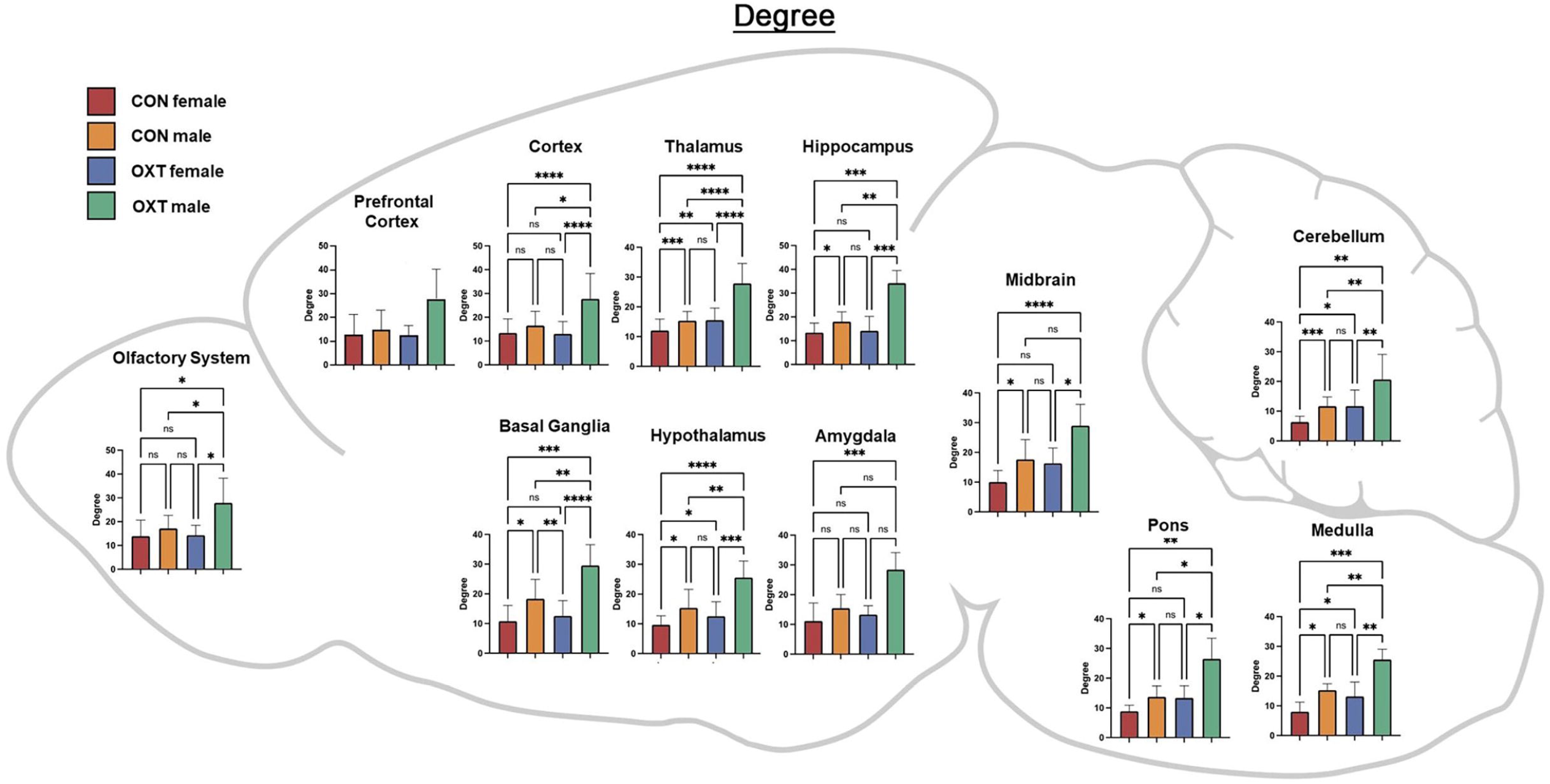
A map of degree averaged over regional clusters by group. Both OXT treatment and male sex increased closeness throughout the brain, with effects most apparent in OXT males. * p < 0.0332, ** p < 0.0021, *** p < 0.0002, **** p <0.0001.

## DISCUSSION

Here we describe how, in prairie voles, exposure to exogenous OXT at birth can impact neurodevelopment in ways that impact neural anatomy and functioning into adulthood. Overall, anatomical differences, while widespread, were generally small, whereas differences in functional connectivity, particularly among OXT-exposed males, were larger. Anatomically, OXT at birth led to a slight reduction in amygdalar volume and OXT males in particular had slightly smaller cortices and slightly larger brainstem / cerebellums (Tables 1-3). OXT at birth led to males resembling females in terms of FA and ADC (Figure 3). However, functionally, OXT males showed marked differences from all other groups. OXT at birth led males to display robustly increased functional connectivity throughout the brain (Figures 4-8). This was particularly the case across the cortex in males in terms of the strength, closeness, and degree of connectivity (Figures 5, 7, and 8). Both the number (Figure 4A) and strength (Figure 4B) of positively correlated connections were greater in males and OXT-exposed animals. This was true both within cluster and, to a lesser extent, between clusters as well (Figure 4C). When these effects were examined regionally, few regions stood out (Figure 5); thus, we view these effects as reflecting broad, brain-wide differences. In terms of graph theoretical analyses of functional connectivity: the basal ganglia stood out for both sex and OXT increasing betweenness, whereas such differences were more widespread for closeness and degree. What does this notably broad increase in functional connectivity mean for the OXT males? We have at present only a few hints.

Interestingly, the observed effects of OXT at birth extended beyond brain regions with dense expression of the OXT receptor. The robust and widespread changes in OXT males’ neural physiology (i.e. functional connectivity) were greater than the changes observed in neuroanatomy (i.e. VBM and DWI). If OXT males are continuously experiencing high levels of communication between brain regions, we would expect that to eventually produce changes in anatomical connectedness. Either our anatomical measures of connectivity were insufficiently sensitive to detect these changes, or the relatively small changes in anatomy we did detect are sufficient to produce functional changes that are comparatively more robust. The functional connectivity scans were undertaken with subjects lightly anesthetized, so it is unlikely that OXT males were responding differentially to the conditions of scanning. Furthermore, we have observed no evidence of stress reactivity being affected by perinatal OXT in our previous studies. OXT acts as a pleiotropic hormone around the time of birth to coordinate the transition from fetal to neonatal life (1), so by affecting neurodevelopmental trajectory, a single OXT exposure could produce such widespread differences across the brain.

We highlight one set of findings in particular because of their relevance to our previous work. As adults, males exposed to OXT via maternal administration at birth had denser OXT receptor distributed along the extent of the agranular insular cortex (14). In the present study, OXT treatment at birth led to the agranular insular cortex registering as a smaller volume in the brains of adult males (Table 3), along with a slight increase in FA (Table 5) and decrease in ADC (Table 8). Moreover, OXT males’ agranular insular cortex had a greater number of functionally connected regions (35 vs. 19 for Control males) and stronger average connectivity among those regions (z-score 3.45 vs. 2.98 for Control males). Changes in the functioning of the agranular insular cortex correspond to the behavioral effects previously observed, such as increased alloparental caregiving (14).

**Table 7.** The list of brain regions that were significantly different in apparent diffusion coefficient (ADC) between Control females and Control males. While widespread, differences were generally small in effect, with males generally showing greater ADC scores.

| Apparent Diffusion Coefficient: Control Females and Males |  |  |  |  |  |  |  |
| --- | --- | --- | --- | --- | --- | --- | --- |
| Brain Area | Females |  |  | Males |  | P val | Effect |
|  | Mean | SD |  | Mean | SD |  |  |
| cuneate nucleus | 1.27 | 0.11 | < | 1.48 | 0.29 | 0.008 | 0.1919 |
| 9th cerebellar lobule | 1.31 | 0.15 | < | 1.52 | 0.28 | 0.009 | 0.18698 |
| tegmental nucleus | 1.29 | 0.15 | < | 1.49 | 0.16 | 0.000 | 0.18492 |
| 7th cerebellar lobule | 1.13 | 0.09 | < | 1.29 | 0.28 | 0.025 | 0.17274 |
| parafascicular thalamic nucleus | 1.24 | 0.11 | < | 1.41 | 0.18 | 0.001 | 0.17138 |
| central medial thalamic nucleus | 1.22 | 0.09 | < | 1.40 | 0.18 | 0.001 | 0.17085 |
| dentate gyrus | 1.40 | 0.15 | < | 1.60 | 0.22 | 0.003 | 0.17017 |
| red nucleus | 1.26 | 0.14 | < | 1.44 | 0.19 | 0.003 | 0.16745 |
| perirhinal ctx | 1.34 | 0.15 | < | 1.52 | 0.24 | 0.009 | 0.16346 |
| habenula nucleus | 1.34 | 0.11 | < | 1.52 | 0.24 | 0.009 | 0.15647 |
| central amygdaloid nucleus | 1.32 | 0.11 | < | 1.49 | 0.18 | 0.002 | 0.1552 |
| ventral tegmental area | 1.45 | 0.23 | < | 1.64 | 0.30 | 0.045 | 0.15392 |
| reticular nucleus | 1.29 | 0.09 | < | 1.46 | 0.18 | 0.002 | 0.15359 |
| caudate putamen (striatum) | 1.23 | 0.09 | < | 1.39 | 0.18 | 0.002 | 0.15305 |
| accumbens core | 1.25 | 0.10 | < | 1.40 | 0.15 | 0.001 | 0.14924 |
| posterior thalamic nucleus | 1.24 | 0.11 | < | 1.39 | 0.17 | 0.004 | 0.14636 |
| globus pallidus | 1.22 | 0.10 | < | 1.37 | 0.17 | 0.004 | 0.14432 |
| zona incerta | 1.32 | 0.16 | < | 1.47 | 0.21 | 0.018 | 0.14322 |
| lateral amygdaloid nucleus | 1.40 | 0.14 | < | 1.56 | 0.20 | 0.008 | 0.14169 |
| temporal ctx | 1.30 | 0.13 | < | 1.45 | 0.21 | 0.016 | 0.14155 |
| median raphe nucleus | 1.33 | 0.16 | < | 1.48 | 0.21 | 0.019 | 0.13934 |
| parabrachial nucleus | 1.31 | 0.14 | < | 1.46 | 0.17 | 0.008 | 0.13793 |
| medial geniculate | 1.26 | 0.12 | < | 1.40 | 0.17 | 0.007 | 0.13656 |
| basal amygdaloid nucleus | 1.31 | 0.17 | < | 1.45 | 0.18 | 0.021 | 0.13487 |
| pontine reticular nucleus oral | 1.34 | 0.21 | < | 1.48 | 0.22 | 0.047 | 0.13411 |
| pretectal nucleus | 1.28 | 0.10 | < | 1.42 | 0.18 | 0.007 | 0.13291 |
| reticular formation | 1.33 | 0.14 | < | 1.47 | 0.19 | 0.015 | 0.13201 |
| claustrum | 1.23 | 0.11 | < | 1.36 | 0.17 | 0.009 | 0.13179 |
| White Matter | 1.37 | 0.11 | < | 1.51 | 0.20 | 0.011 | 0.13133 |
| reuniens nucleus | 1.28 | 0.13 | < | 1.42 | 0.18 | 0.015 | 0.1309 |
| ventral thalamic nuclei | 1.28 | 0.11 | < | 1.41 | 0.18 | 0.010 | 0.13063 |
| subiculum | 1.40 | 0.16 | < | 1.55 | 0.20 | 0.021 | 0.12898 |
| dorsal raphe | 1.35 | 0.14 | < | 1.49 | 0.20 | 0.020 | 0.12782 |
| lateral septal nucleus | 1.45 | 0.12 | < | 1.60 | 0.18 | 0.008 | 0.12336 |
| auditory ctx | 1.29 | 0.12 | < | 1.42 | 0.19 | 0.023 | 0.12044 |
| periaqueductal gray | 1.41 | 0.11 | < | 1.55 | 0.22 | 0.027 | 0.11978 |
| lateral posterior thalamic nucleus | 1.30 | 0.10 | < | 1.43 | 0.17 | 0.012 | 0.11941 |
| CA3 | 1.39 | 0.13 | < | 1.53 | 0.21 | 0.028 | 0.11874 |
| posterior hypothalamic area | 1.32 | 0.17 | < | 1.44 | 0.18 | 0.039 | 0.1185 |
| infralimbic ctx | 1.48 | 0.16 | < | 1.62 | 0.18 | 0.018 | 0.1179 |
| CA1 | 1.40 | 0.12 | < | 1.53 | 0.19 | 0.017 | 0.11749 |
| bed nucleus stria terminalis | 1.30 | 0.13 | < | 1.43 | 0.17 | 0.019 | 0.11585 |
| orbital ctx | 1.36 | 0.14 | < | 1.48 | 0.21 | 0.046 | 0.11332 |
| solitary tract nucleus | 1.30 | 0.13 | < | 1.42 | 0.20 | 0.044 | 0.11314 |
| paraventricular nuclus | 1.37 | 0.15 | < | 1.49 | 0.16 | 0.022 | 0.11212 |
| medial septum | 1.34 | 0.14 | < | 1.46 | 0.14 | 0.022 | 0.10701 |
| lateral geniculate | 1.31 | 0.11 | < | 1.42 | 0.18 | 0.034 | 0.10652 |
| extended amygdala | 1.33 | 0.15 | < | 1.44 | 0.17 | 0.043 | 0.10648 |

**Table 8.** The list of brain regions that were significantly different in apparent diffusion coefficient (ADC) between OXT males and Control males. While widespread, differences were generally small in effect, with Control males generally showing greater ADC scores. Note: there were no regions where OXT females and Control females differed in terms of ADC.

| Apparent Diffusion Coefficient: Control Males vs Oxytocin Males |  |  |  |  |  |  |  |
| --- | --- | --- | --- | --- | --- | --- | --- |
| Brain Area | Males Control |  |  | Males Oxytocin |  | P val | Effect |
|  | Mean | SD |  | Mean | SD |  |  |
| periaqueductal gray | 1.55 | 0.22 | > | 1.31 | 0.14 | 0.000 | 0.25266 |
| tegmental nucleus | 1.49 | 0.16 | > | 1.26 | 0.13 | 0.000 | 0.24327 |
| vestibular nucleus | 1.48 | 0.16 | > | 1.26 | 0.15 | 0.000 | 0.22849 |
| dorsal raphe | 1.49 | 0.20 | > | 1.28 | 0.14 | 0.000 | 0.22449 |
| parafascicular thalamic nucleus | 1.41 | 0.18 | > | 1.22 | 0.12 | 0.000 | 0.22109 |
| reuniens nucleus | 1.42 | 0.18 | > | 1.22 | 0.14 | 0.000 | 0.22122 |
| paraventricular nuclus | 1.49 | 0.16 | > | 1.29 | 0.13 | 0.000 | 0.21074 |
| posterior thalamic nucleus | 1.39 | 0.17 | > | 1.21 | 0.13 | 0.000 | 0.20552 |
| superior colliculus | 1.45 | 0.19 | > | 1.26 | 0.13 | 0.000 | 0.20326 |
| ventral thalamic nuclei | 1.41 | 0.18 | > | 1.23 | 0.14 | 0.000 | 0.19862 |
| CA3 | 1.53 | 0.21 | > | 1.33 | 0.15 | 0.000 | 0.20163 |
| central amygdaloid nucleus | 1.49 | 0.18 | > | 1.31 | 0.14 | 0.000 | 0.19427 |
| central medial thalamic nucleus | 1.40 | 0.18 | > | 1.22 | 0.12 | 0.000 | 0.19809 |
| lateral amygdaloid nucleus | 1.56 | 0.20 | > | 1.39 | 0.13 | 0.000 | 0.17413 |
| medial dorsal thalamic nucleus | 1.40 | 0.19 | > | 1.22 | 0.14 | 0.000 | 0.20071 |
| pretectal nucleus | 1.42 | 0.18 | > | 1.25 | 0.12 | 0.000 | 0.18295 |
| secondary somaotsensory ctx | 1.42 | 0.17 | > | 1.27 | 0.13 | 0.000 | 0.16995 |
| medial geniculate | 1.40 | 0.17 | > | 1.24 | 0.14 | 0.000 | 0.18186 |
| bed nucleus stria terminalis | 1.43 | 0.17 | > | 1.26 | 0.13 | 0.000 | 0.18427 |
| accumbens core | 1.40 | 0.15 | > | 1.24 | 0.12 | 0.000 | 0.17873 |
| CA1 | 1.53 | 0.19 | > | 1.36 | 0.15 | 0.000 | 0.17772 |
| lateral dorsal thalamic nucleus | 1.48 | 0.18 | > | 1.32 | 0.13 | 0.000 | 0.16654 |
| parabrachial nucleus | 1.46 | 0.17 | > | 1.30 | 0.16 | 0.000 | 0.16313 |
| lateral posterior thalamic nucleus | 1.43 | 0.17 | > | 1.26 | 0.14 | 0.000 | 0.17875 |
| red nucleus | 1.44 | 0.19 | > | 1.27 | 0.14 | 0.000 | 0.18789 |
| primary somatosensory ctx | 1.55 | 0.21 | > | 1.39 | 0.17 | 0.001 | 0.15046 |
| reticular nucleus | 1.46 | 0.18 | > | 1.29 | 0.13 | 0.001 | 0.17881 |
| inferior colliculus | 1.50 | 0.21 | > | 1.30 | 0.14 | 0.001 | 0.20488 |
| habenula nucleus | 1.52 | 0.24 | > | 1.35 | 0.19 | 0.001 | 0.16646 |
| medial septum | 1.46 | 0.14 | > | 1.29 | 0.13 | 0.001 | 0.17305 |
| paraventricular nucleus | 1.40 | 0.19 | > | 1.23 | 0.17 | 0.001 | 0.18977 |
| White Matter | 1.51 | 0.20 | > | 1.35 | 0.15 | 0.001 | 0.16027 |
| lateral geniculate | 1.42 | 0.18 | > | 1.25 | 0.13 | 0.001 | 0.17768 |
| 2nd cerebellar lobule | 1.47 | 0.20 | > | 1.30 | 0.21 | 0.001 | 0.17439 |
| caudate putamen (striatum) | 1.39 | 0.18 | > | 1.23 | 0.12 | 0.001 | 0.16938 |
| claustrum | 1.36 | 0.17 | > | 1.21 | 0.11 | 0.001 | 0.16714 |
| anterior thalamic nuclei | 1.40 | 0.18 | > | 1.24 | 0.15 | 0.001 | 0.17436 |
| median raphe nucleus | 1.48 | 0.21 | > | 1.33 | 0.17 | 0.001 | 0.14888 |
| reticular formation | 1.47 | 0.19 | > | 1.31 | 0.19 | 0.001 | 0.16269 |
| accumbens shell | 1.43 | 0.16 | > | 1.26 | 0.14 | 0.002 | 0.18139 |
| auditory ctx | 1.42 | 0.19 | > | 1.27 | 0.16 | 0.002 | 0.1609 |
| globus pallidus | 1.37 | 0.17 | > | 1.21 | 0.14 | 0.002 | 0.17193 |
| orbital ctx | 1.48 | 0.21 | > | 1.31 | 0.14 | 0.003 | 0.17342 |
| infralimbic ctx | 1.62 | 0.18 | > | 1.44 | 0.16 | 0.003 | 0.16252 |
| 7th cerebellar lobule | 1.29 | 0.28 | > | 1.08 | 0.18 | 0.003 | 0.26364 |
| visual 1 ctx | 1.50 | 0.25 | > | 1.34 | 0.19 | 0.003 | 0.15891 |
| anterior olfactory nucleus | 1.57 | 0.30 | > | 1.35 | 0.22 | 0.003 | 0.21393 |
| 10th cerebellar lobule | 1.62 | 0.25 | > | 1.43 | 0.17 | 0.003 | 0.18096 |
| dentate gyrus | 1.60 | 0.22 | > | 1.41 | 0.19 | 0.003 | 0.18057 |
| agranular insular ctx | 1.45 | 0.17 | > | 1.30 | 0.23 | 0.004 | 0.1574 |
| 5th cerebellar lobule | 1.25 | 0.24 | > | 1.07 | 0.19 | 0.004 | 0.22931 |
| extended amygdala | 1.44 | 0.17 | > | 1.29 | 0.17 | 0.004 | 0.15247 |
| 3rd cerebellar lobule | 1.30 | 0.22 | > | 1.13 | 0.21 | 0.004 | 0.20307 |
| endopiriform nucleus | 1.40 | 0.18 | > | 1.26 | 0.18 | 0.005 | 0.15393 |
| 6th cerebellar lobule | 1.24 | 0.23 | > | 1.03 | 0.18 | 0.005 | 0.26201 |
| parasubiculum | 1.57 | 0.28 | > | 1.42 | 0.21 | 0.005 | 0.14979 |
| subiculum | 1.55 | 0.20 | > | 1.40 | 0.24 | 0.007 | 0.13718 |
| frontal association ctx | 1.58 | 0.24 | > | 1.43 | 0.19 | 0.008 | 0.15083 |
| zona incerta | 1.47 | 0.21 | > | 1.31 | 0.22 | 0.011 | 0.17364 |
| temporal ctx | 1.45 | 0.21 | > | 1.30 | 0.22 | 0.012 | 0.15813 |
| lateral septal nucleus | 1.60 | 0.18 | > | 1.48 | 0.12 | 0.012 | 0.10938 |
| primary motor ctx | 1.60 | 0.27 | > | 1.49 | 0.19 | 0.014 | 0.09447 |
| visual 2 ctx | 1.66 | 0.31 | > | 1.52 | 0.20 | 0.015 | 0.12248 |
| ventral pallidum | 1.47 | 0.18 | > | 1.32 | 0.18 | 0.015 | 0.15104 |
| pontine reticular nucleus oral | 1.48 | 0.22 | > | 1.36 | 0.27 | 0.016 | 0.12224 |
| solitary tract nucleus | 1.42 | 0.20 | > | 1.28 | 0.21 | 0.016 | 0.14862 |
| retrosplenial ctx | 1.77 | 0.33 | > | 1.64 | 0.22 | 0.016 | 0.10686 |
| basal amygdaloid nucleus | 1.45 | 0.18 | > | 1.30 | 0.23 | 0.022 | 0.16251 |
| anterior hypothalamic area | 1.55 | 0.23 | > | 1.39 | 0.27 | 0.023 | 0.1591 |
| 9th cerebellar lobule | 1.52 | 0.28 | > | 1.33 | 0.19 | 0.024 | 0.18827 |
| posterior hypothalamic area | 1.44 | 0.18 | > | 1.28 | 0.16 | 0.024 | 0.17083 |
| medial preoptic area | 1.57 | 0.25 | > | 1.41 | 0.23 | 0.024 | 0.15719 |
| diagonal band of Broca | 1.53 | 0.22 | > | 1.40 | 0.21 | 0.025 | 0.13414 |
| pontine reticular nucleus caudal | 1.49 | 0.22 | > | 1.38 | 0.34 | 0.025 | 0.1033 |
| perirhinal ctx | 1.52 | 0.24 | > | 1.36 | 0.31 | 0.028 | 0.16618 |
| anterior cingulate ctx | 1.75 | 0.24 | > | 1.58 | 0.20 | 0.028 | 0.14404 |
| crus ansiform lobule | 1.36 | 0.24 | > | 1.23 | 0.23 | 0.029 | 0.1512 |
| 8th cerebellar lobule | 1.38 | 0.29 | > | 1.23 | 0.24 | 0.034 | 0.1625 |
| gigantocellular reticular nucleus | 1.53 | 0.26 | > | 1.43 | 0.38 | 0.036 | 0.10108 |
| granular cell layer olfactory bulb | 1.56 | 0.30 | > | 1.40 | 0.16 | 0.040 | 0.16061 |
| cuneate nucleus | 1.48 | 0.29 | > | 1.34 | 0.25 | 0.042 | 0.14091 |
| dorsal medial nucleus | 1.64 | 0.25 | > | 1.51 | 0.20 | 0.046 | 0.11454 |
| medial amygdaloid nucleus | 1.68 | 0.25 | > | 1.52 | 0.34 | 0.049 | 0.13551 |

One pattern generally observed in the brains of humans with autism spectrum disorders is diminished long-distance functional connectivity and increased short-distance functional connectivity (27,28). We observed that both male sex and OXT treatment at birth led to increased intra-cluster connectivity (analogous to increased short-distance hyperconnectivity), but also to increased inter-cluster connectivity, though to a lesser extent. Thus, the present results do not entirely resemble the equivalent effects seen in humans with autism spectrum disorders. While some epidemiological studies have suggested a link between autism spectrum disorders and perinatal oxytocin exposure (11), residual confounding by either genetic or environmental vulnerabilities, could explain this apparent association. Indeed, our previous work, we observed a broadly gregarious phenotype in OXT-exposed voles (14); if such results translated to humans, they would support the contention that underlying vulnerabilities bring on both a need for OXT in the mother and susceptibility to autism spectrum disorders in the child. While our previous work found behavioral differences in both males and females exposed to OXT, which is somewhat in contrast to the present study’s findings, we also found males to be much more affected by perinatal OXT in terms of neuroanatomy, which was assessed as the density of OXT and AVP receptors -and that finding matches those of the present study. Numerous previous studies have found sex-, dose-, and region-dependent effects of early life oxytocin manipulation (9,29).

There have been very few studies on the impact of birth interventions and subsequent brain development (1,3). In the realm of neuroanatomy, Deoni and colleagues recently reported that CS results in smaller brain volumes in neonatal mouse pups (30). However, no such findings have been observed in human children or infants (31). In terms of VBM and functional connectivity in the present study, OXT led males toward a more masculinized phenotype. In terms of ADC and FA, however, OXT led males toward a more feminized phenotype. Further work is needed to reveal the meanings of these differences.

This study is not without limitations. Firstly, because the Control group did not receive a vehicle treatment, the effects of injection were not adequately controlled for, which introduced a stress confound of indeterminate magnitude. In a small validation study, adult offspring of saline-treated dams (n = 3 female, 5 male adult offspring) were not found to have meaningful differences in DWI values compared to the un-treated Control animals of the present study (n = 17 female, 19 male adult offspring).

Collectively, these results support the contention that the perinatal brain is sensitive to OXT administered indirectly to the pregnant female. As obstetric care continues to use OXT for labor induction / augmentation in the majority of births in the U.S. (4,5), a more complete understanding of the neurodevelopmental effects of OXT exposure at delivery is imperative. Furthermore, as delayed cord clamping becomes standard practice (32), this will extend both the duration and dose of potential OXT exposure by the neonate, since the vast majority of deliveries now use OXT during the third stage of labor to prevent postpartum hemorrhage (33,34). Because the perinatal period is a sensitive period for brain development in terms of OXT exposure, these practices deserve further investigation.

## Supplementary Figure Legends

Table S1. The list of brain regions and their corresponding regional cluster to which they were classified.

Figure S1. The changes in brain volumes between sexes are not consistent, i.e. in some brain areas females are greater (black sign) and others less than males (red sign). The location of these different brain areas affected by OT at birth when comparing females and males can be seen in the probability heat maps in Fig 1. Brain regions that are greater in volume in females than males are organized around the olfactory system (A. glomerular layer; B. anterior olfactory n; B-D. piriform cortices) and prefrontal/limbic cortical areas (B. motor, insular; C. anterior cingulate; D. retrosplenial). Areas that are larger in males vs females are primarily associated with brainstem/cerebellum (E. reticulotegmental n., raphe, pontine reticular n; F. olivary n.; G. vestibular n., gigantocellularis, paraflocculus, F. cuneate n. 7th lobule, medullary reticular n.).

## Notes

### Competing Interest Statement

The authors have declared no competing interest.

### Summary of Updates

The manuscript text has been edited to include more Discussion as well as a revised Results section to reflect the inclusion of new Figures.

